# Single-cell Senseless protein analysis reveals metastable states during the transition to a sensory organ fate

**DOI:** 10.1101/2021.12.20.473498

**Authors:** Ritika Giri, Shannon Brady, Dimitrios K. Papadopoulos, Richard W. Carthew

**Affiliations:** Department of Molecular Biosciences, Northwestern University, Evanston IL 60208; NSF-Simons Center for Quantitative Biology, Northwestern University, Evanston IL 60208; Center for Molecular Medicine (CMM), Department of Clinical Neuroscience, Karolinska Institute, 17176 Stockholm, Sweden; Department of Biology, University of Crete, Voutes University Campus, 70013 Heraklion, Crete, Greece

**Keywords:** Drosophila, Senseless, sensory neuron development

## Abstract

Cell fate decisions can be envisioned as bifurcating dynamical systems, and the decision that *Drosophila* cells make during sensory organ differentiation has been described as such. We extended these studies by focusing on the Senseless protein, which orchestrates sensory cell fate transitions. Wing cells contain intermediate Senseless numbers prior to their fate transition, after which they express much greater numbers of Senseless molecules as they differentiate. However, the dynamics are inconsistent with it being a simple bistable system. Cells with intermediate Senseless are best modeled as residing in four discrete states, each with a distinct protein number and occupying a specific region of the tissue. Although the states are stable over time, the number of molecules in each state vary with time. The fold-change in molecule number between adjacent states is invariant and robust to absolute protein number variation. Thus, cells transitioning to sensory fates exhibit metastability with relativistic properties.

## INTRODUCTION

Animal development requires the progressive restriction of cells into distinct fate lineages. Before reaching their terminal fate, most cells transition through a number of stable lineage states, such that the fate potential of a cell is progressively restricted with each transition. C.H. Waddington famously compared the path that a cell takes towards its ultimate fate to a ball rolling down a contoured landscape and coming to rest at a terminal (Waddington, 1957).

Attractor states have long been proposed to represent distinct stable cell states, with transitions between attractor states being modeled as bifurcation events (Corson and Siggia, 2012; Huang et al., 2005). Waddington’s landscape imagines there to be pitchfork bifurcations, where one state bifurcates into two stable states. With such bifurcations, the dynamical system is initially in a single stable steady state. Beyond the bifurcation, the original state remains a steady state but becomes unstable. Two new stable steady states appear, and these are symmetrical. In spite of the graphical similarity between Waddington’s metaphor and pitchfork bifurcations in dynamical systems (Marco et al., 2014), there are other means by which cells could make fate choices over time. Saddle-node or fold bifurcations create new stable steady states somewhere distant from the initial steady state, which ceases to exist. Unlike pitchfork bifurcations, the initial steady state is always unstable and is intermediate between the two stable states. Therefore, a dynamical system does not remain for a significant amount of time in this intermediate state but rapidly moves into a new stable state.

It is non-trivial to distinguish whether cell fate transitions should be modeled as pitchfork or saddle-node bifurcations (Freedman et al., 2021). Both could be invoked to explain the behavior of molecular determinants of cell fates (Marco et al., 2014; Rand et al., 2021). Fate transitions often involve the intermediate expression of a molecular determinant, which is then interpreted in a dichotomous manner by individual cells, to adopt divergent fates (Levine et al., 2013). For example, mouse neural progenitor cells co-express fluctuating levels of three transcription factors, Ascl1/Mash1, Hes1 and Olig2, which promote fate choices to become neurons, astrocytes, and oligodendrocytes, respectively (Imayoshi et al., 2013). Subsequently, each differentiated lineage is characterized by high sustained expression of only one of the factors.

A different example is seen in developing structures that give rise to the adult *Drosophila* body. Sensory bristles on the adult epidermis derive from cells called sensory organ precursors (SOPs). Each SOP cell emerges from a group of cells expressing intermediate levels of proneural transcription factors from the *Achaete-Scute* (*Ac-Sc*) gene complex and the *senseless* (*sens*) gene (Garcia-Bellido and de Celis, 2009; Jafar-Nejad et al., 2003; Jafar-Nejad et al., 2006). The SOP is selected from the group by the action of Notch-mediated lateral inhibition, leading to sustained high expression of Sens and Ac-Sc proteins by a positive feedback mechanism (Nolo et al., 2000, 2001). In contrast, cells not selected for SOP fate downregulate Sens and Ac-Sc expression, and become epidermal cells. Mathematical modeling of the system as a saddle-node bifurcation has successfully described these dynamics, according to which an initial unstable state of intermediate proneural gene expression resolves into two stable states with high and low proneural expression (Corson et al., 2017).

When considering cell fate transitions as described above, it is generally assumed that the initial steady state has a fixed average expression level of fate determinants, with noise creating continuous variation in levels between cells. However, mammalian pluripotent embryonic stem (ES) cells can express fate determinants heterogeneously such that they resemble discrete states of expression. It is thought that ES cells can reversibly transit between these discrete states, thereby becoming temporarily biased to adopt specific differentiated fates (Klein et al., 2015). These states have been termed metastable, akin to the definition in physics of an energy state with a longer lifetime than that generated by random fluctuation but with a shorter lifetime than the ground state. From a dynamical perspective, the definition of metastability varies according to the approaches dealing with the issue (Kelso, 2012). For example, metastability has been described as a system that explores its state space on different time scales. On fast time scales, transitions happen within a single subregion of state space, and on slow time scales they occur between different subregions. Metastability has also been described as a regime near a saddle-node bifurcation in which stable and unstable states no longer exist but attraction remains to where those fixed points used to be. However, it is unclear if metastable cell states are a general feature of developmental fate transitions.

In order to explore the phenomenon of metastable cell-states during developmental transitions, we investigated the well-characterized process of SOP selection. We chose to focus on development of chemosensory organs located at the anterior margin of the adult wing (Fig. 1A). The wing develops from an elliptically shaped columnar epithelium known as the wing pouch, which grows in size during the larval life stages (Fig. 1B). SOP cells that give rise to the chemosensory organs first appear at the transition between the larval and pupal stages of life. The SOPs develop in two periodic rows flanking the midline that bisects the dorsal and ventral halves of the wing pouch (Fig. 1C). This can be microscopically visualized by the high level of Sens expression in these cells (Fig. 1D). Some time later, each SOP divides twice, and the four descendants differentiate into a single sensory organ.

**Figure 1.**
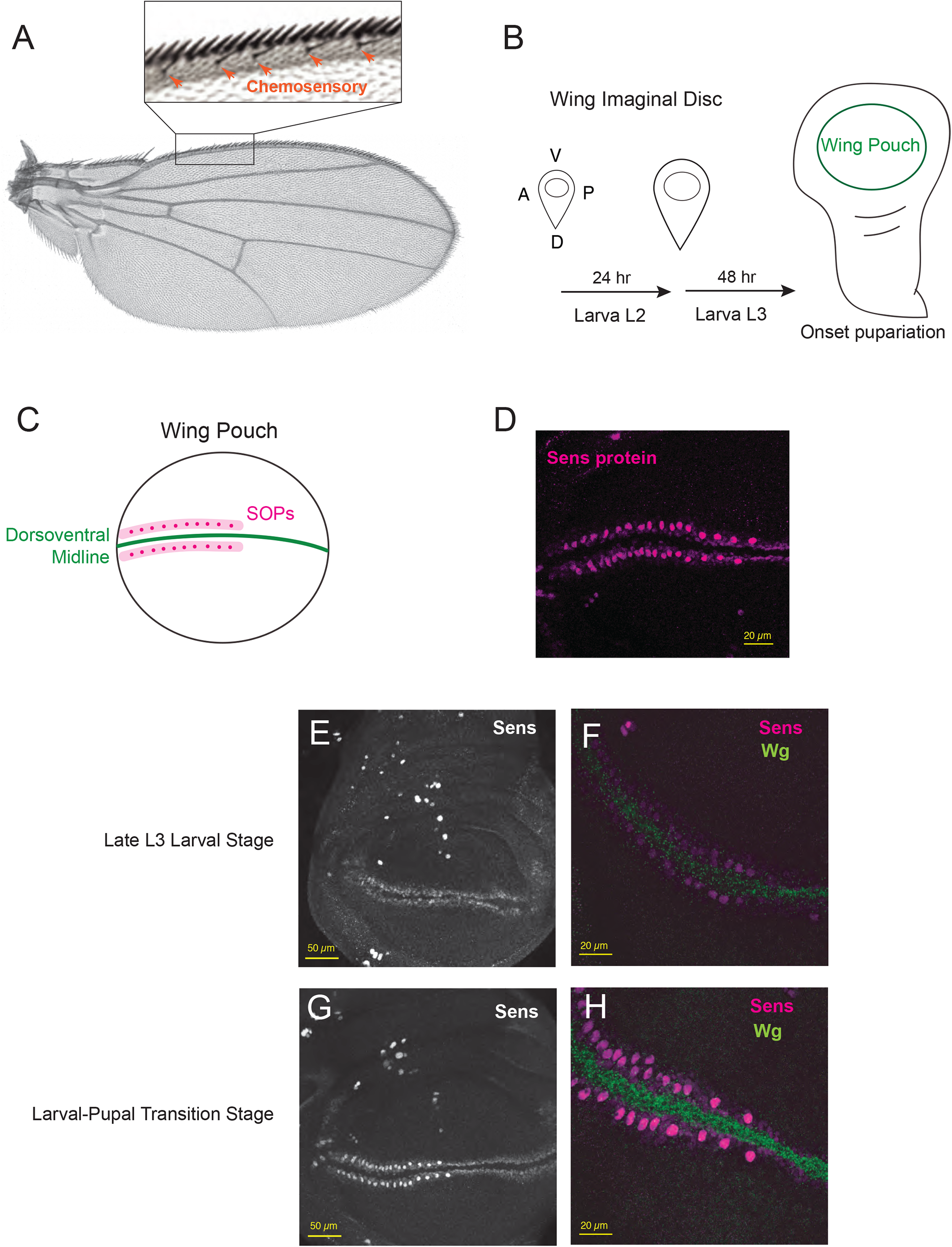
Sensory organ fate selection at the developing *Drosophila* wing margin. (A) An adult wing with the anterior margin on top. Inset shows a magnified view of the margin with slender chemosensory bristles marked. Stout mechanosensory bristles occupy the extreme margin edge. (B) Schematic of the wing imaginal disc as it grows during larval stages until pupation. Embedded in the disc is the wing pouch as indicated. A, anterior; P, posterior; D, dorsal; V, ventral. (C) Schematic of Sens expression in the wing pouch at the larval-pupal transition stage. (D) Sens protein fluorescence in a wing pouch. Note the periodic bright spots of fluorescence denoting SOPs. (E,F) In the late larval stage, two stripes of Sens protein (white in E and purple in F) are induced by Wg protein (green in F). Panel E is a lower magnification view of the pouch. (G,H) In the larval-pupal transition stage, the two stripes of Sens protein (white in G and purple in H) evolve to have within them two periodic rows of SOPs. Panel G is a lower magnification view of the pouch.

Sens expression is first detected during the larval stage prior to detection of the SOP cells (Fig. 1E). It is initially expressed at low levels within two stripes that are 3 – 6 cells in width and flank the dorsal-ventral midline. Expression is induced by the Wnt signaling protein Wingless (Wg), which is secreted by midline cells and whose expression increases over time (Fig. 1F,H) (Jafar-Nejad et al., 2006; Zecca et al., 1996). A subset of Sens-positive cells then dramatically increase Sens expression and become SOPs (Fig. 1G,H); these are typically located in the center of each stripe. This selection process is mediated by Delta-Notch mediated lateral inhibition (Hinz et al., 1994). Cells expressing higher levels of Sens express Delta ligand more strongly (Nolo et al., 2001). This leads to stronger Notch receptor activation in neighbors, resulting in inhibition of Sens expression and down-regulation of Delta in the neighbors (Jafar-Nejad et al., 2003). These neighbors are then less effective at inhibiting Sens in the stronger expressing cell, leading to even stronger expression. Since Sens protein can activate *Ac-Sc* expression and its own expression (Acar et al., 2006; Jafar-Nejad et al., 2003; Nolo et al., 2000), the Delta-Notch lateral inhibitory process is further amplified. Thus, small initial differences in the levels of Sens protein between neighboring cells evolve into large differences (Fig. S1A,B). The system is thought to have bistability, such that intermediate levels of Sens are unstable, and cells transit to either high Sens (SOPs) or low Sens (epidermal) stable states (Fig. S1C).

## RESULTS

An elegant mathematical model was formulated to describe SOP pattern formation of the dorsal thorax (Corson et al., 2017). Dynamics of cell fate patterning were encoded by modeling a 2D lattice of cells. Cells expressed a fate potential that varied over simulation time according to the dynamics of signaling, and they exhibited bistability under intermediate signal levels. Cells then adopted a SOP fate potential (maximal) in the absence of inhibitory signals and an epidermal fate potential (minimal) under strong inhibition. Bistability arose due to the non-linear behavior of the inhibitory signal and feed-back amplification of the activating signal. A periodic pattern of SOP cells emerged along multiple parallel rows of cells, much like the observed SOP pattern in the developing thorax (Corson et al., 2017).

We adapted this model to mimic development of the wing margin SOPs. We modified the model to contain a 1D array of cells in order to resemble a single stripe of proneural cells beside the wing margin (see Methods). As model simulations proceeded over time, fate potential progressively increased in all cells until an intermediate fate potential was reached. This transient steady state represents an unstable one, since the system has a saddle-node bifurcation (Fig. 2A). At this point, the system evolved such that some cells adopted a SOP fate potential and some cells adopted an epidermal fate potential (Fig. 2B). The 1D array of cells self-organized into a pattern of periodically spaced SOPs (Fig. 2C). Although our model simulations predicted fate dynamics that could be tested in live wing pouch tissue, the conditions to culture wing pouches *ex vivo* and maintain their intact development are not yet achievable (Handke et al., 2014; Zartman et al., 2013). Thus we could not directly observe dynamics by live imaging. Instead, wing imaginal discs were dissected out and fixed at different developmental stages.

**Figure 2.**
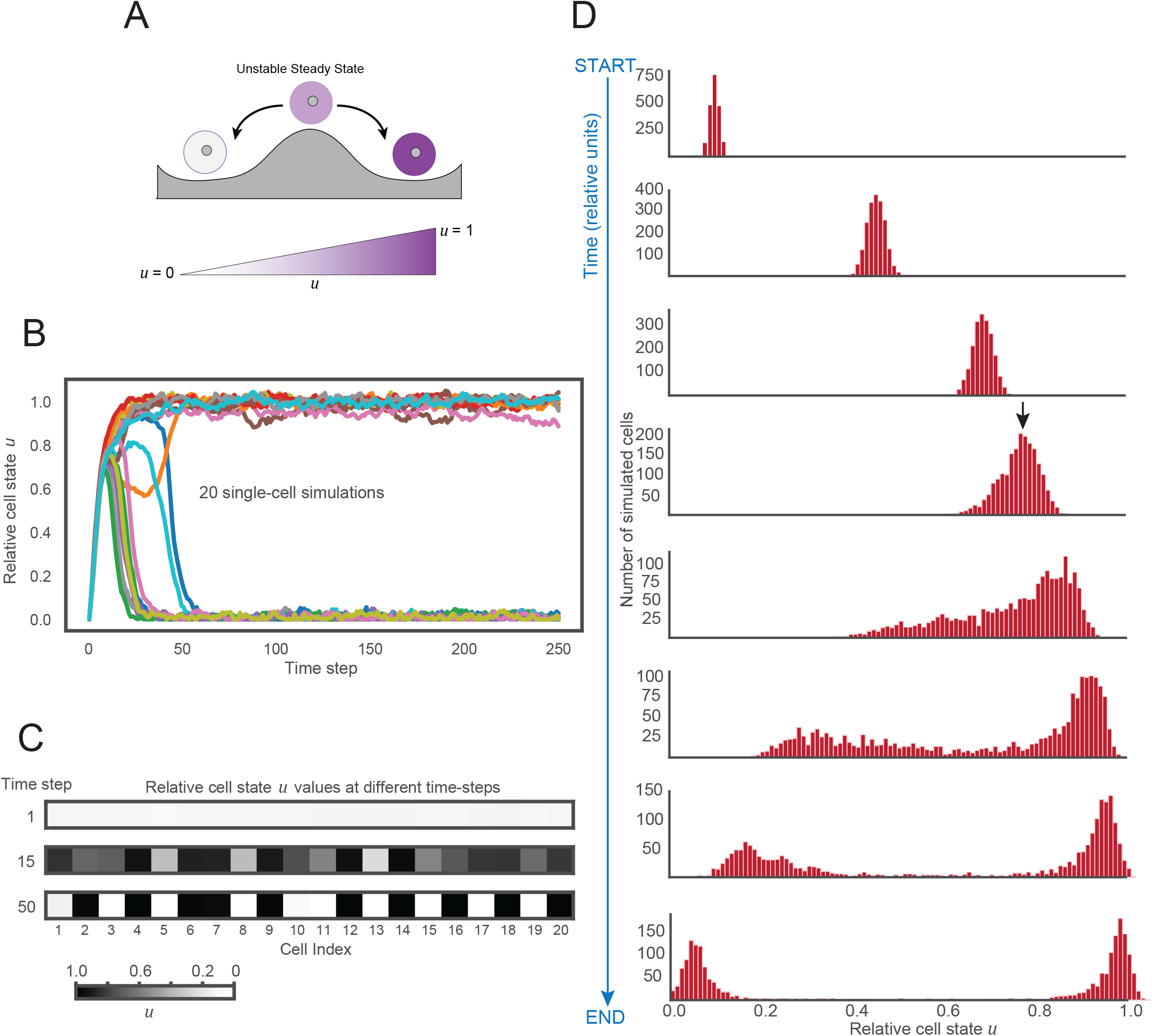
A mathematical model for Sens pattern formation. (A) The SOP vs epidermal fate choice was modeled as a system with bistable dynamics, and the variable *u* representing SOP fate-potential (Corson et al., 2017). The model was implemented such that cells were arranged in a 1D array with periodic boundary conditions. Each cell in the 1D cell array reached an unstable steady state of intermediate value for SOP fate-potential *u*. Cells then bifurcated to zero or maximal values of *u*, corresponding to stable steady states. (B) Single-cell traces for *u* as simulations evolved over time. A total of 20 simulated cells are shown comprising a single simulation run. (C) A representative 1D array of 20 cells shown as they varied in *u* over simulation time. Note the “errors” in periodic patterning between cells 6-7 and 10-11. Such patterning errors are generated due to the noise term present in the model. The noise parameter was set to 1% relative to *u*. A lower noise parameter value resulted in fewer “errors”. (D) The distribution of the cell state variable *u* in model simulations over time. A total of 100 simulations with 20 cells each (total 2,000 cells) were run with identical parameters. At discrete time steps, simulations were tallied for their values of *u*. The arrow marks the time at which the unstable steady state begins to bifurcate. In the model, Sens protein level is assumed synonymous with the cell state variable *u*.

To make a prediction from the model that we could experimentally test, we simulated the cell distributions of fate-potential at different time points during pattern resolution. As expected, this revealed that at early time points, fate-potential displayed a unimodal distribution. The mean of this unimodal distribution steadily increased until a certain point, at which it split into a bimodal distribution (Fig. 2D). This provided us with a testable prediction of the model since we could approximate fate-potential by measuring the levels of Sens protein in many single cells from wing pouches fixed at different time points of development.

Such an approximation assumes that Sens protein abundance is a reasonable proxy for fate potential in the wing margin. This assumption is motivated by several lines of evidence. Clonal analysis indicates that in the absence of Sens, Ac-Sc expression fails to be upregulated in wing margin SOP cells (Jafar-Nejad et al., 2003). This is markedly different from the thoracic SOPs where Sens does not direct fate selection, but rather the later division and differentiation of microchaete progeny cells (Jafar-Nejad et al., 2006). Moreover, Sens misexpression has been shown to be sufficient to induce cells to become SOPs (Nolo et al., 2000). Further, our previous work exploring stochastic noise in Sens expression revealed that modest increases in the noise of the Sens protein expression led to the formation of ectopic wing chemosensory organs (Giri et al., 2020). Finally, Sens dynamics are well-suited to bistable behavior since it molecularly acts as a transcription activator at high concentrations, but as a transcription repressor at low concentrations. This orchestrates SOP selection by feedback amplification of Sens’s own transcription (Jafar-Nejad et al., 2003; Powell et al., 2012).

### A mixture model describes Sens protein distribution

To measure Sens protein levels in single cells, we employed a genomic *sens* transgene tagged at the N-terminus with sfGFP. The transgene fully recapitulates the endogenous *sens gene* expression pattern, and fully rescues the developmental defects of *sens* mutant alleles (Giri et al., 2020). All experiments were performed with two copies of the transgene in a *sens* mutant background.

Single-cell Sens levels were quantified using a previously described automated image analysis pipeline (Giri et al., 2020). Wing pouches from transgenic animals were fixed and imaged with a confocal microscope. Single cells were identified by computational image segmentation, and Sens was quantified as the average sfGFP fluorescence signal in each segmented cell. Relative fluorescence signals were converted to absolute sfGFP-Sens molecule numbers using a calibration factor that was empirically determined by Fluorescence Correlation Spectroscopy (FCS). FCS relies on recording fluorescence intensity fluctuations from a sub-femtoliter confocal volume with high, sub-microsecond temporal resolution reaching single-photon detection efficiencies (Giri et al., 2020) (for details, see Methods).

To ensure that the recorded FCS fluctuations are generated by molecular movement, we performed additional experiments. We first fitted FCS measurements with a two-component diffusion model. Then, we ascertained that the sfGFP-Sens fast and slow characteristic decay times, indicating freely diffusing and chromatin-bound molecules, respectively, remained stable as a function of the total sfGFP-Sens concentration. This indicated that the observed fluorescence intensity fluctuations were caused by molecular movement (i.e., diffusion) of sfGFP-Sens molecules through the detection volume. We conducted similar measurements with Sens tagged with a different fluorescent protein (mCherry-Sens), and observed comparable behaviors to sfGFP-Sens. Additionally, we observed that the mobility measured by FCS was largely unaffected by changes in nuclear sfGFP-Sens and mCherry-Sens concentrations (Fig S2B,C). Further, we established that the shape of the autocorrelation curves recorded at different stages of larval and pupal wing disc development remained unaltered. This indicated that the fractions of the freely diffusing and chromatin-bound Sens molecules, as well as their mobilities, do not change substantially among different developmental stages of the wing discs (Fig. S2D). To preclude a possible contribution of energy transfer between sfGFP-Sens and mCherry-Sens, we examined the concentration of sfGFP-Sens as a function of the total mCherry-Sens concentration and found no correlation between the two (Fig. S2A). Moreover, we examined how the total number of sfGFP-Sens molecules and its characteristic times τ_1_ and τ_2_ (derived by fitting) change as a function of the size of the fluorescence detection volume. Both characteristic times and the number of sfGFP-Sens molecules increased linearly with the detection volume (Fig. S2E–G). This indicated that molecular motion underlies the observed fluorescence intensity fluctuations and provided no indication of anomalous diffusion. Finally, we performed consecutive FCS measurements in the same cells, in which we varied the laser excitation intensity of sfGFP-Sens molecules 2-, 5-, and 10-fold. We observed that the photon counts per molecule and second of mCherry-Sens molecules remained largely unaltered. Thus, we obtained no evidence for the existence of Förster Resonance Energy Transfer between sfGFP-Sens and mCherry-Sens, indicating that energy transfer processes contribute poorly, if at all, to the fluctuations. We conclude that the molecular count factor derived previously (Giri et al., 2020) using sfGFP-Sens/mCherry-Sens cells could be faithfully used to calculate absolute Sens concentrations in homozygous sfGFP-Sens cells.

For each experimental condition in our study, between 6,000 and 12,000 Sens-positive cells were segmented and analyzed. Wing pouches were dissected from late third instar larvae ∼ 1 - 2 hours before SOPs are first detected. Several wing pouches were subjected to the automated analysis pipeline and pooled to generate a high-resolution expression profile for Sens (Fig. 3A). Sens-positive cells displayed a wide range of protein numbers. However, 90% of cells expressed fewer than 2,000 molecules per cell, and the majority of those contained fewer than 200 molecules per cell. We then dissected wing pouches from animals undergoing the larval-pupal transition, when the first SOPs have formed. There was a near two-fold increase in Sens numbers per cell relative to the earlier time-point (Fig. 3B). The median number of 188 molecules per cell in larval cells shifted to 374 molecules per cell in the larval-pupal cells. The difference in Sens numbers over time was statistically significant and corresponded to an increase of 186 molecules (95% confidence interval 175-197 molecules). Since SOP cells were visibly apparent at the larval-pupal transition, we counted Sens protein number in these cells and found them to contain from 1,500 to over 10,000 molecules per cell.

**Figure 3.**
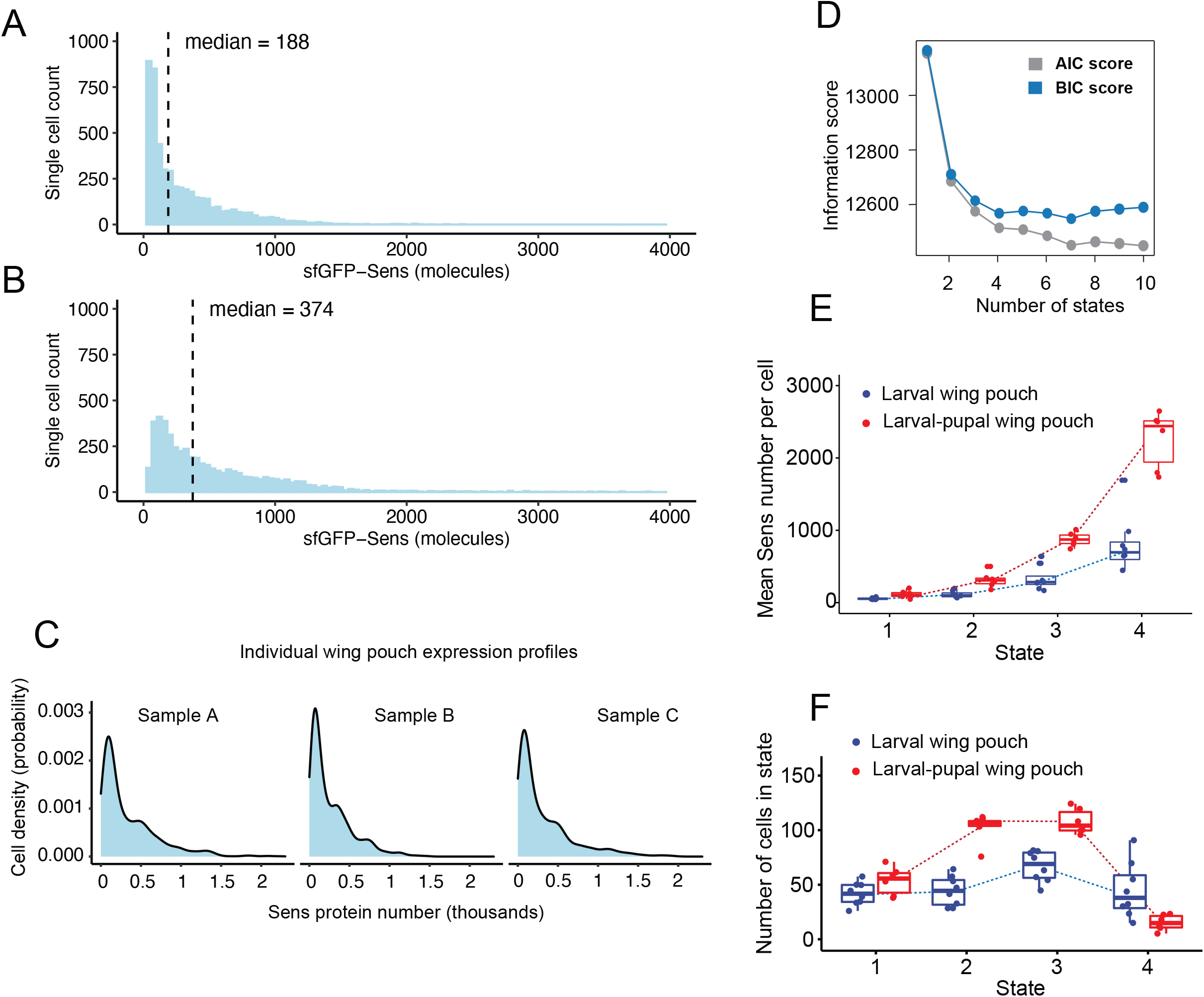
Sens-positive proneural cells display four-state multimodality. (A,B) A histogram of sfGFP-Sens protein number per cell for 5,000 randomly sampled cells is shown in each panel. Cells were sampled at the late larval stage (A) or the larval-pupal transition stage (B). The x-axis is truncated at 4,000 molecules for visualization, however cellular Sens expression can reach up to 10,000 molecules (C) Sens expression profiles (density plots) derived from three representative wing pouches collected at the larval stage. (D) AIC and BIC scores were independently derived for Gaussian mixture models of the Sens data from an individual wing pouch. The AIC and BIC assess the relative quality of statistical models built with different values of the parameter *k*, which defines the number of states within a parent distribution. Models with a *k* value that generates the lowest AIC and BIC scores are deemed to be best-fit, in that they are best at describing the data without overfitting by having higher values of *k*. (E) Mean Sens protein number for cells in the four states. Each dot represents an estimate of the mean derived from one wing pouch sample. Multiple samples were analyzed for each state and condition. Wing pouches are from the larval (blue) and larval-pupal (red) stages of development. Boxplots show the median and interquartile range (25 – 75%), and whiskers represent the 1.5x interquartile range. The difference in Sens number between larval and larval-pupal stages for each state were all statistically significant (*p* < 0.02, Wilcoxon test). (F) Estimated number of cells within each state for the four states. Each dot represents an estimate derived from one wing pouch sample. Wing pouches are from the larval (blue) and larval-pupal (red) stages of development. Boxplots show the median and interquartile range (25 – 75%), and whiskers represent the 1.5x interquartile range.. The difference in cell number between larval and larval-pupal stages for state 1 was not significant (*p* = 0.081) but for states 2,3, and 4, were all statistically significant (*p* < 0.005, Wilcoxon test).

We focused our analysis on cells with intermediate Sens levels since these cells were predicted to be approaching the unstable steady state. Within these cells, there appeared to be a multimodal distribution of Sens protein number, with a single major peak and a number of smaller peaks with higher Sens number (Figs. 3C and S3). In order to rigorously assess the composition of the Sens-positive cell population, Sens distributions obtained from individual wing pouches were subjected to mixture model analysis. A mixture model is a probabilistic model that predicts the number of discrete states present in an overall set of observations, without mapping individual data points to a specific state. We fit a Gaussian mixture model to the Sens expression data. Model quality was determined by calculating the Akaike Information Criterion (AIC) and Bayesian Information Criterion (BIC) scores for models with different numbers of states (Claeskens and Hjort, 2008). A model with four discrete states was repeatedly found to best-fit the distributions observed in replicate wing pouches from both the larval and larval-pupal stages (Fig. 3D, Fig S4). Information scores from mixture models with different numbers of components is shown for a representative wing pouch from the larval-pupal stage (Fig. 3D). This analysis suggests that cells expressing intermediate levels of Sens do not show a simple pattern of variation around a single mean, as predicted by the dynamical model (Fig. 2D), but rather there is unanticipated heterogeneity in their expression.

Each cell state in the Gaussian mixture model is defined by three parameters: the mean Sens protein number, the standard deviation around the mean, and the fractional representation of the state in the total cell population. For instance, a cell state with a mean expression of 100 molecules, standard deviation of 20 molecules, and fractional proportion of 0.3 would indicate that 30% of all Sens-positive cells are in a state centered at 100 #20 Sens molecules. We labelled the different Sens states from 1 to 4, with state 1 having the lowest number of Sens molecules per cell and state 4 having the highest (Table 1). For both the early and late stages, there appeared to be a non-linear increase in average Sens number when comparing one state to the next (Fig. 3E, Fig. S8). When comparing early and late developmental stages, the average Sens number differed by two-fold between states with the same label (Fig. 3E, Fig. S8). However, it is important to note that the similar labeling of states at different stages does not necessarily imply that they are the same. Cells labeled state 1 at the early stage are not necessarily in the same internal state as cells labeled state 1 at the late stage.

**TABLE 1.**
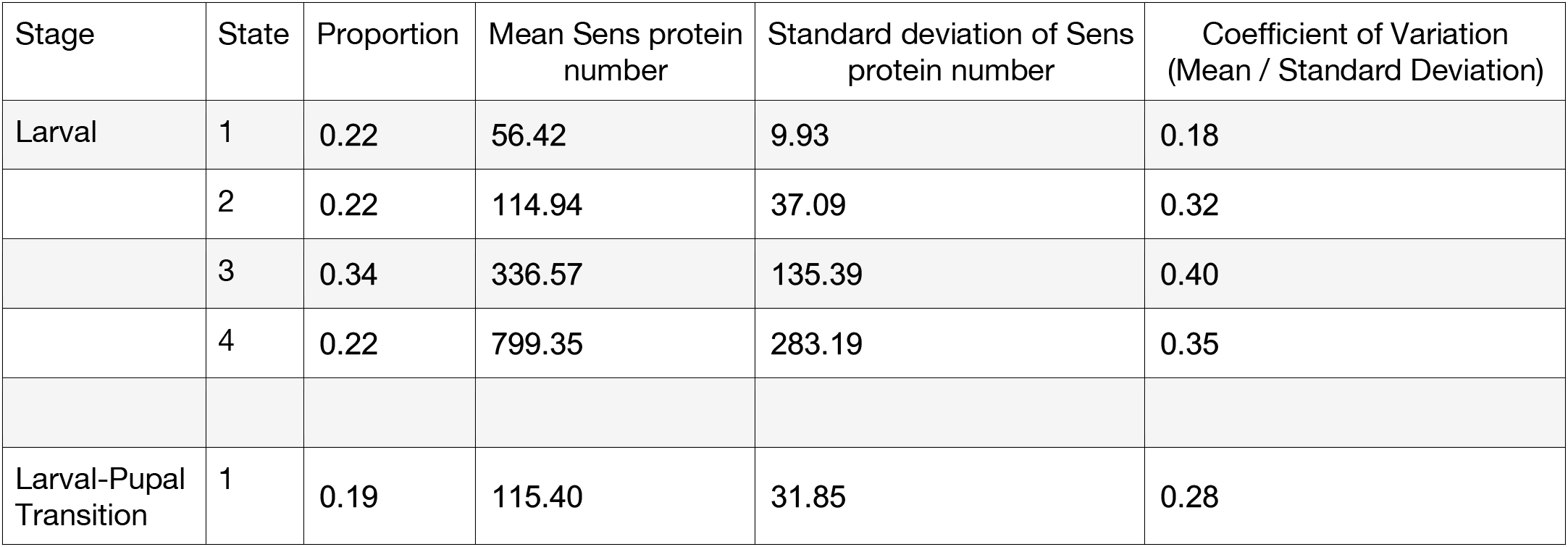
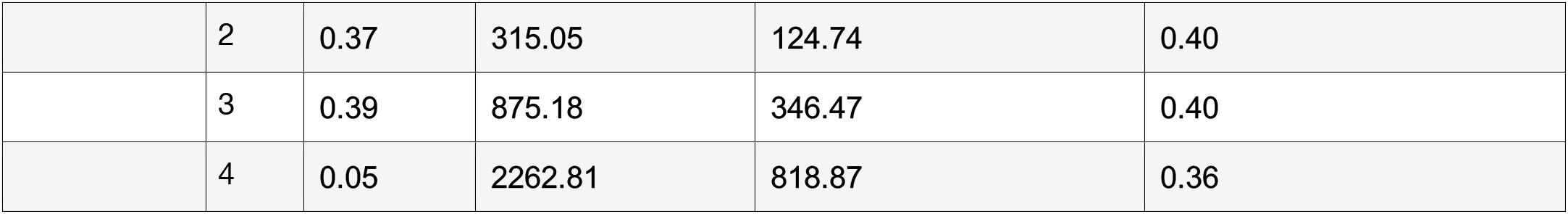
State values from mixture modeling individual wing pouches from n > 6 wing pouches per stage. Note the difference in means between adjacent states is greater than the sum of their standard deviations, suggesting the states are well separated in the models.

We converted the fractional proportion of cells in a given state to the absolute number of cells in that state. Conversion was performed in order to compare stages, since the total number of Sens-positive cells in the imaged regions increased from 198 #41 in larval wing pouches to 279 # 21 in in larval-pupal pouches. At the larval stage, the imaged region in the pouch contained approximately 50 cells in each of the four states (Fig. 3F). At the later larval-pupal stage, there remained 50 cells in state 1, but there were more cells in states 2 and 3, while fewer cells were found to be in state 4 (Fig. 3F). This observation did not fit with expectations of the dynamical model, in which cells bifurcate towards low and high states as time proceeds (Fig. 2D). If that were true, we would have observed more cells in the lowest and highest states of 1 and 4. Instead, more cells were detected in the two intermediate states 2 and 3.

### Cells in different states occupy overlapping regions in the wing pouch

Wg molecules activate *sens* transcription on either side of the dorsal-ventral midline up to eight cell diameters away (Eivers et al., 2009; Giri et al., 2020). Since Wg concentration decays with distance from the midline (Alexandre et al., 2014), it is possible that cells at different distances receive different doses of the activating signal. If Sens expression is proportional to the signal dose, then the resulting distribution would resemble an *n*-state mixture, where *n* distinct doses of Wg are interpreted as *n* cell states.

Therefore, we examined the relationship between the distance of a cell from the Wg source, and its Sens expression level. The midline was manually delineated, and the shortest Euclidean distance from each cell centroid to the midline was calculated (Fig. S5). As expected, Sens-positive cells were located no further than eight cell-diameters from the midline. However, Sens protein numbers were weakly correlated with distance, with correlation coefficients ranging from 0.25 to 0.35 for individual samples (Fig. 4A). At both early and late stages, cells close to the midline displayed a wide range of Sens protein number. As distance from the midline increased, the range of Sens number narrowed, and cells expressed fewer Sens molecules.

**Figure 4.**
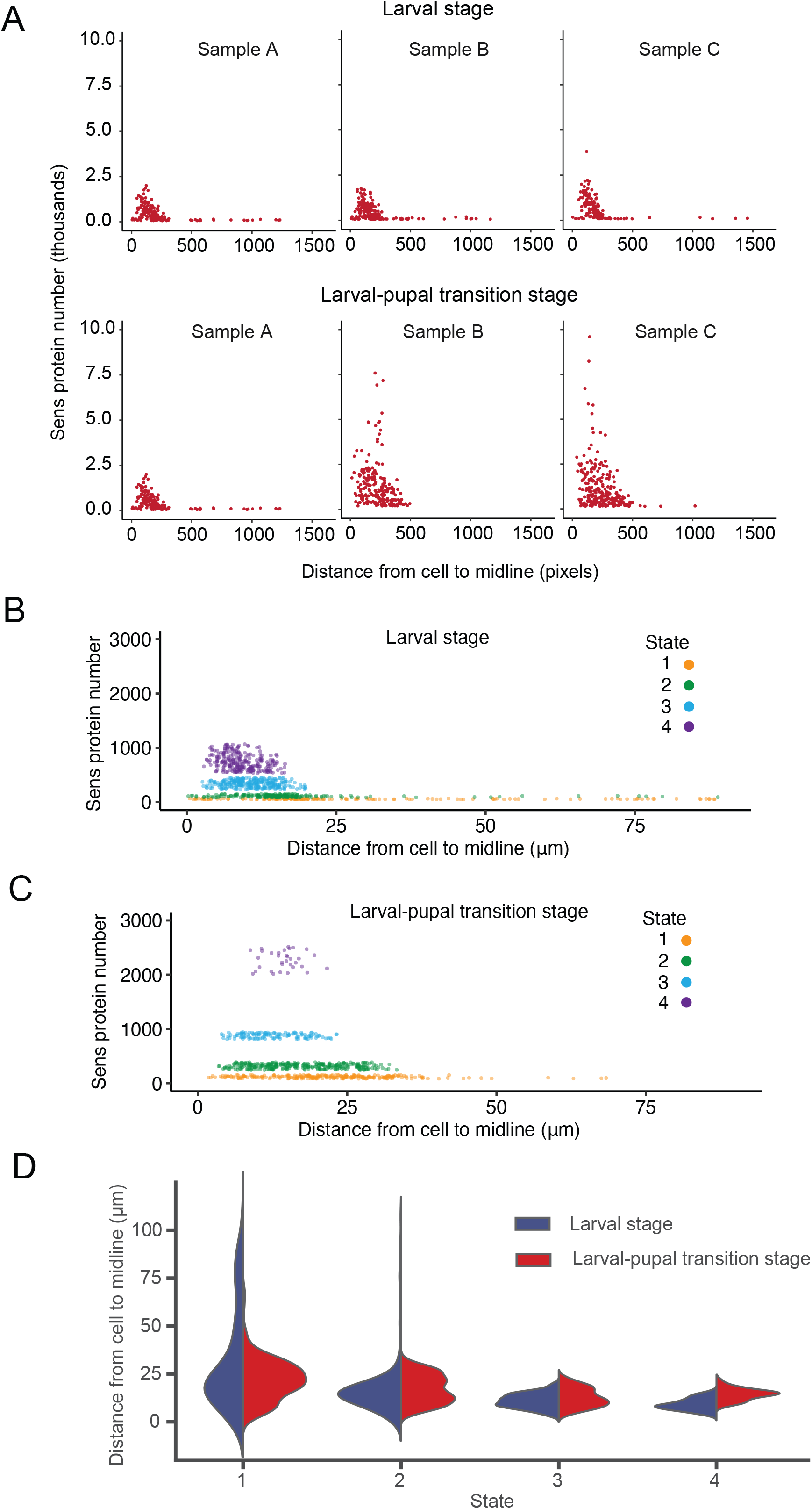
Cells in Sens expression states are patterned within each proneural stripe. (A) Sens expression level is weakly correlated with distance from the dorsoventral midline. Single cells from three representative wing pouches at the larval stage (top) and larval-pupal transition stage (bottom) are plotted for their distance to the midline on the x-axis and Sens protein number on the y-axis. (B,C) Single cells were classified into one of the states according to their Sens protein number relative to the mean Sens number in states. Cells are color coded according to state classification as shown, and plotted as a function of their distance to the midline on the x-axis and Sens protein number on the y-axis. Shown are cells analyzed from one wing pouch taken at the larval stage (B) or larval-pupal transition stage (C). (D) Half-violin plots of cells’ distance to the dorsoventral midline according to their state classification. Cells from larval stage and larval-pupal transition stage are plotted together according to their state classification.

To ascertain where cells in a given state are located, we classified cells according to their likely state. This was done by assigning cells with Sens levels within one standard deviation from the mean value of a given state. Each cell’s distance from the dorso-ventral midline was then calculated as before. At the larval stage, cells had a nested spatial distribution within the proneural stripe (Fig. 4B, Fig. S9). Cells in state 1 were distributed over the greatest distance, followed by cells in state 2, state 3 and state 4, each with progressively more restricted spatial distributions. In a zone located 5 to 15 μm from the dorso-ventral midline, cells in all four states coexisted (Fig. 4B, Fig. S9). At the larval-pupal transition, the nested spatial distribution of states was also observed (Fig. 4C, Fig. S9). However, the distributions were narrower; the distal boundaries of cells in states 1 and 2 were much closer to the dorso-ventral midline. The proximal boundary of cells in state 4 was more distant from the dorso-ventral midline, and the distal boundary also appeared to have shifted. When we mapped visible SOPs to these distributions, the SOPs were located in the central zone within the proneural stripe, in which cells in all four states resided. The nested distribution of states within each proneural stripe suggests that Wg dose alone does not determine Sens state.

The spatial position of cells in various states was conserved between larval and larval-pupal stages (Fig. 4D). There was large overlap in position between cells labelled state 1 at the early stage and cells labelled state 1 at the late stage, and so on. Cells in the wing pouch do not move relative to one another at this period of wing development. Hence, the results suggest that individual states are maintained over time, i.e. cells labelled state 4 at the early stage are probably in the same internal state as those labelled state 4 at the later stage. If a given state is maintained over time, this would suggest that between the early and late stages, some cells transitioned between states. The reason for this supposition is that some regions with cells in a given state at the early stage no longer had such cells later (Fig. 4D). For example, state 1 cells very distal from the midline at the larval stage likely became Sens-negative by the pupal transition. State 2 cells distal from the midline at the larval stage either became Sens-negative or transitioned to state 1. The overall implication is that while internal states are maintained over time, some cells transition into or out of particular states over time.

### States are robust to perturbation of absolute Sens number

If internal states are maintained over time, they are nevertheless dynamic in terms of Sens molecule number. There is a two-fold increase in Sens number within cells of a given state from early to late stages. There is also a change in the number of cells in a given state over time. It is therefore possible that an increase in Sens number alters the likelihood that cells remain in a particular state. To test the idea, Sens expression was genetically perturbed, and the resulting distribution was analyzed to determine cell-state properties. Sens expression was perturbed by ablating miR-9a regulation of *sens*. MicroRNAs are non-coding RNA molecules that bind to target mRNAs in a sequence-specific manner and modestly repress protein output (Carthew and Sontheimer, 2009). We and others had shown that miR-9a inhibits Sens protein output by binding to two sites in the *sens* mRNA 3’UTR (Cassidy et al., 2013; Giri et al., 2020; Li et al., 2006). To test the effect of miR-9a regulation on cell state properties, the miR-9a binding sites in the *sens* transgene were combinatorially mutated to generate a series of *sens* alleles. Alleles were landed in the same genomic location as the wild-type *sens* transgene. All transgenes rescued *sens* mutant phenotypes and recapitulated the endogenous *sens* expression pattern (Fig. S6A).

The single binding site mutants *sens-m1* and *sens-m2* displayed only modest changes in Sens expression (Fig. S6B). When both binding sites were mutated in *sens-m1m2*, there was a significant increase in Sens expression (Figs. 5A and S6C). This indicates that the two binding sites are functionally redundant *in vivo*, such that a single site can confer near-complete repression. With loss of both binding sites, median single-cell Sens levels increased from 374 molecules to 616 molecules per cell, a 1.65-fold increase (Fig. 5A). Increased expression was a direct consequence of loss of miR-9a binding, since the expression profiles did not significantly differ in a *mir-9a* mutant background (Fig. S6D).

**Figure 5.**
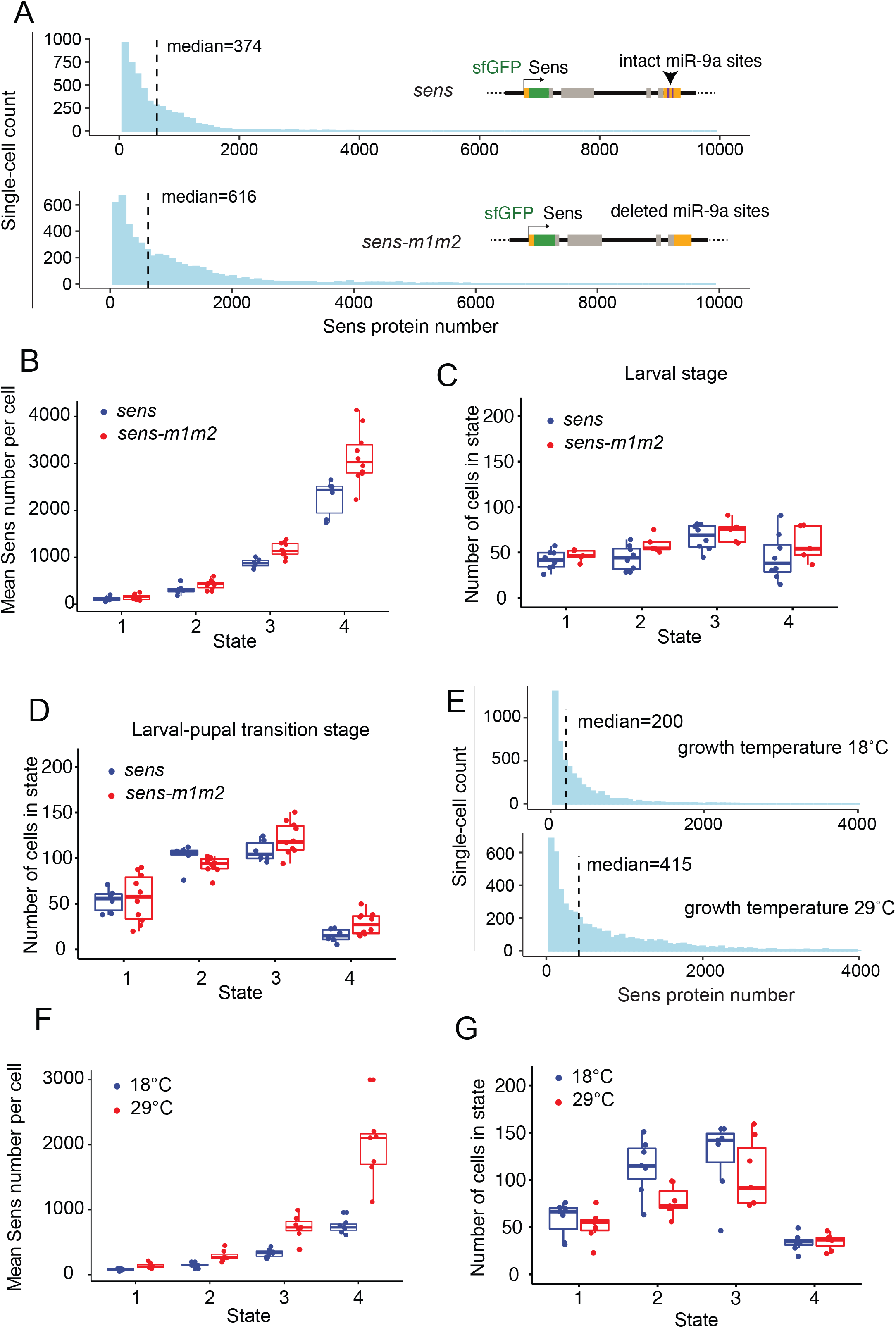
Perturbation of Sens number per cell has little effect on cell-state properties. (A) Histograms of sfGFP-Sens protein number in 5,000 randomly sampled cells are shown. Dotted lines represent the median numbers for the populations. Cells were collected from larval-pupal cells expressing wild-type *sens* (top) or the miR-9a binding-site mutant allele *sens-m1m2* (bottom). (B) The mean Sens protein number for cells in the four states. Each dot represents a mean estimate of protein number derived from one larval-pupal stage wing pouch. Multiple samples were analyzed for each condition. Wing pouches are from wildtype *sens* (blue) and *sens-m1m2* (red) alleles. Boxplots show the median and interquartile range (25 – 75%), and whiskers represent the 1.5x interquartile range. The differences in Sens number between *sens* and *sens-m1m2* for states 1 and 2 were not significant (*p* > 0.10, Wilcoxon test) but for states 3 and 4 they were significant (*p* < 0.003, Wilcoxon test). (C,D) Estimated number of cells within each state for the four states. Each dot represents an estimate derived from one wing pouch. Wing pouches are from wildtype *sens* (blue) and *sens-m1m2* (red) alleles. Boxplots show the median and interquartile range (25 – 75%), and whiskers represent the 1.5x interquartile range. Cells were sampled at the larval (C) and larval-pupal transition (D) stages. The differences in cell number between *sens* and *sens-m1m2* for each state in (C) were not significant (*p* > 0.09, Wilcoxon test). The differences in cell number between *sens* and *sens-m1m2* for states 1, 3, and 4 in (D) were not significant (*p* > 0.07, Wilcoxon test) but for state 2 it was significant (*p* = 0.02, Wilcoxon test). (E) Histograms of sfGFP-Sens protein number in 5,000 randomly sampled cells are shown. Dotted lines represent the median numbers for the populations. Cells were collected from larval-pupal cells grown at 18°C (top) or 29°C (bottom). (F) Sens protein number for cells in the four states. Each dot represents a mean estimate of protein number derived from one larval-pupal stage wing pouch. Wing pouches are from animals grown at 18°C (blue) and 29°C (red). Boxplots show the median and interquartile range (25 – 75%), and whiskers represent the 1.5x interquartile range. The differences in Sens number between 18°C and 29°C for each state were all significant (*p* < 0.0025, Wilcoxon test). (G) Estimated number of cells within each state for the four states. Each dot represents an estimate derived from one larval-pupal stage wing pouch. Wing pouches are from animals grown at 18°C (blue) and 29°C (red). Boxplots show the median and interquartile range (25 – 75%), and whiskers represent the 1.5x interquartile range. The differences in cell number between 18°C and 29°C for states 1, 3, and 4 were not significant (*p* > 0.30, Wilcoxon test) but for state 2 it was significant (*p* = 0.04, Wilcoxon test).

Cell populations from the *sens-m1m2* wing pouches were then subject to mixture model analysis. Again, four states were determined to be the best-fit model for the mutant expression profile (Fig S4). The average Sens molecule number in each state was uniformly higher when miR-9a regulation was ablated, suggesting that miR-9a regulates expression in a state-independent manner (Fig. 5B, Fig. S8B). Importantly, there was little effect of the *sens-m1m2* mutant on the number of cells in each state at both larval and larval-pupal stages (Fig. 5C,D). These results suggest that the cell states are robust against loss of miR-9a regulation of protein output. Hence, the developmental change in cell-state population size seen in wildtype is not simply due to the increase in Sens protein number over that time period. If it had been, then cell-state population sizes would have differed between the wildtype and *sens* mutant at the same developmental stage, which was not observed.

To further test the insensitivity of cell states to absolute Sens numbers, we raised animals at different temperatures. The previous experiments had all been conducted at a room temperature of 22 - 24°C. Two other temperatures, 18°C and 29°C, were chosen to assess cell state properties at the larval-pupal transition. At both 18°C and 29°C, the Sens expression pattern appeared normal (Fig. S7A). However, median Sens number was two-fold higher when growth temperature was increased from 18°C to 29°C (Fig. 5E).

Sens-positive cell populations from 18°C and 29°C were subjected to mixed model analysis to determine if cell state characteristics were altered. Again, a mixed model with four states was the best-fit whether at 18 or 29°C. The average Sens levels uniformly increased for all four states from 18°C to 29°C (Fig. 5F, Fig. S8C). However, no significant difference was observed in cell-state population size between the two temperatures (Fig. 5G). As a final test, animals with different *sens* alleles were subjected to temperature variation. Compared to wildtype *sens* animals raised at 18°C, *sens-m1m2* animals raised at 29°C showed a four-fold increase in average Sens protein number for all states (Fig. S7B, Fig. S8C). However, there was little change in cell-state population size (Fig. S7C). Altogether, these results indicate that the relative number of cells in each state is robust against variation in Sens number.

### States differ by a fixed ratio of Sens protein numbers per cell

We next analyzed the relative levels of Sens in states by calculating the ratio of average Sens molecule number between adjacent states. The ratios of Sens in state 2 vs state 1, state 3 vs state 2, and state 4 vs state 3 were strikingly similar to one another (Fig. 6A), in spite of the large differences in absolute Sens protein number between adjacent states (Figs. 3E and 5B,F). Remarkably, these ratios remained invariant when measured in cells at the larval stage, at the larval-pupal stage, when raised at 18°C or 29°C, and when miR-9a regulation of *sens* was nullified (Fig. 6A). There were no significant differences between the ratios under all of these conditions (*p* > 0.130 for all Z-tests). Indeed the relativistic nature of the states can be best visualized when Sens numbers in each state are plotted on logarithmic scales. Regardless of the type of *sens* allele, developmental stage or growth temperature, we observed a log-linear trend of increasing Sens from the lowest to highest cell states (Fig. S8). This result indicates that the proneural system elicits fold-changes in Sens number when cells transition to particular states. If this interpretation is correct, then states are not determined so much by an absolute number of Sens molecules as by a relative number.

**Figure 6.**
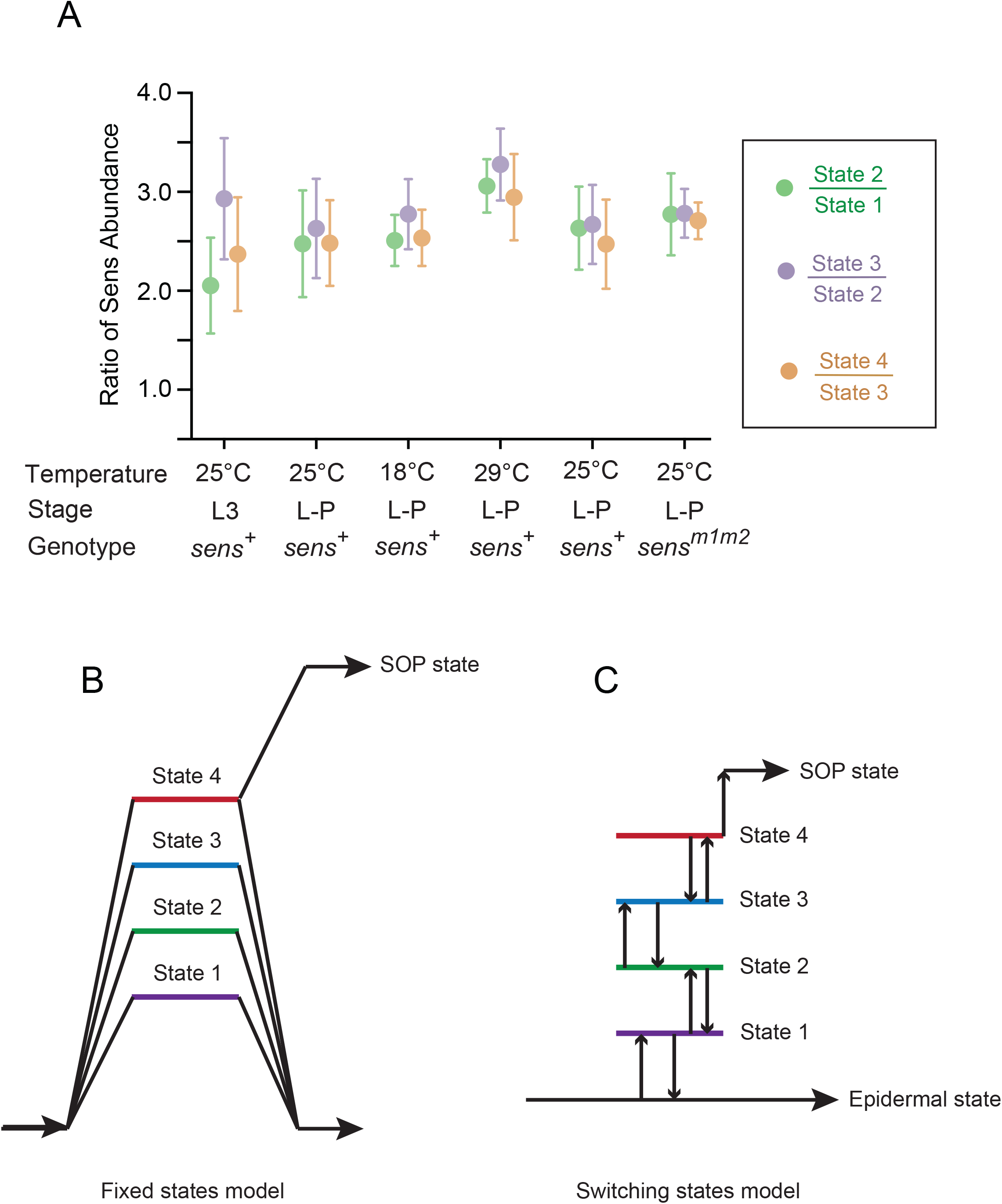
Relative ratio of Sens number and state models. (A) Shown is the ratio between the mean number of Sens molecules per cell measured in adjacent states: state 2 vs 1; state 3 vs 2; state 4 vs 3. Each datapoint corresponds to this ratio of means estimated from samples taken from a particular combination of experimental conditions. These conditions are growth temperature, developmental stage (L3: larval stage; L-P: larval-pupal transition stage), and the *sens* transgene allele (*sens^+^*: wildtype *sfGFP-sens*; *sens^m1m2^*: *sfGFP-sens* with miR-9a binding sites mutated). Error bars represent the standard error of the mean. Replicate wings for each sample were analyzed to derive these ratios. The number of replicates per sample ranged from 7 to 10 wings. There are no statistically significant differences between the state ratios within any of the six experimental conditions (*p* > 0.130 for all Z-tests). Although the state ratios observed at 29°C are modestly higher than for the other conditions, there are no significant differences when compared to the ratios observed at 25°C (*p* > 0.148 for all Z-tests). (B,C) Two alternative models for how cells transition to and from metastable states. (B) Cells transition from a ground state to any one of four metastable states. They then either relax back to the ground state or transition to an SOP state with some likelihood, in this example with only state 4 being permissive for transition to the SOP state. Other scenarios are also possible. (C) Cells transition from a ground state to a particular metastable state, and following this, they transition to and from all of the other metastable states with some likelihood. In this example, only state 1 cells can relax back to the ground state and only state 4 cells can transition to an SOP state, but many other scenarios are equally plausible.

## DISCUSSION

A simple saddle-node bifurcation model of SOP selection at the wing margin predicts that before the bifurcation, Sens levels become unimodally distributed around an intermediate unstable steady state. Once selected, the model predicts SOP-fated precursors continuously increase Sens to its maximal value, while epidermal-fated precursors continuously reduce Sens to its minimal value. However, single-cell protein quantitation during development *in vivo* reveals an unanticipated distribution of cells into four discrete subpopulations expressing low-to-intermediate levels of Sens. These subpopulations are maintained in the face of differences in developmental stage, growth temperature, and miR-9a regulation, indicating that the four-state distribution is a stable property of Sens expression in proneural cells of the wing margin.

The existence of stable subpopulations suggests the system has metastability. There are two distinct ways in which the system could create this metastability. One, cells could begin to express Sens and then continuously increase Sens number until cells reach any one of the four possible states (Fig. 6B). The likelihood a cell adopts a particular state might depend on variables such as its distance from the midline. However, once a cell adopts that particular state, its state is fixed such that it does not transition to one of the other three states, but it either relaxes to being Sens-negative or it transitions to the SOP fate. The likelihood a cell in a particular state can become an SOP might vary with its state identity, i.e. have zero likelihood if it is in state 1, 2, and 3 but some positive likelihood if it is in state 4. This scenario would be consistent with the fact that all SOPs emerge only in the narrow zone where cells in state 4 reside.

A second way in which the system could have metastability would be that cells can transition between different metastable states with some switching probability dependent on a variable such as distance from the midline (Fig. 6C). For example, a cell in state 1 could switch to state 2 and vice versa. Transitions could conceivably be limited to adjacent states or might include switches to non-adjacent states, i.e. state 1 to state 3. If the system operates this way, then the transition of a competent proneural cell to its SOP fate is not a continuous process but requires multiple transitions between metastable states before the SOP fate is adopted. Such metastable systems have been observed in other developmental contexts, and have been shown to have functional roles (Antolovic et al., 2019; Chang et al., 2008; Klein et al., 2015; Olsson et al., 2016).

This raises the question as to what gives rise to these discrete states? One possibility is that burst-like transcription of genes generates discrete pulses of protein number that accumulate over time. Transcription initiation is rarely a continuous process for genes, even those that are constitutive. Short periods of time in which multiple transcripts are initiated are interspersed by periods of time in which no initiation occurs. If the average timescale between bursts is long enough, then discrete bursts of mRNA number lead to multimodality in protein number (Thomas et al., 2014). Indeed, modeling of *sens* transcription in the wing margin suggests that in lowly expressing cells, timescales of the on and off periods are approximately equivalent (Bakker et al., 2020; Giri et al., 2020). These studies also indicate that an average of four *sens* mRNAs are generated per burst, and each mRNA is translated on average into ten Sens protein molecules (Giri et al., 2020). Moreover, since Sens protein turnover is on a much longer time-scale, with a half-life of five hours (Giri et al., 2020), then a transcription burst would result in the accumulation of ∼ 40 Sens protein molecules. Since the average Sens protein number in state 1 cells at the larval stage is 56 and at the larval-pupal stage is 109, it is plausible that state 1 represents cells after one or two transcriptional bursts of *sens*. State 2 could also conceivably be created by bursting kinetics. The average Sens number in state 2 cells at the larval stage is 115 and at the larval-pupal stage is 270, which would correspond to two and five transcription bursts, respectively. Burst size remains constant in cells experiencing different levels of Sens expression (Bakker et al., 2020).

However, a model in which transcription bursts generate discretization of all four states has a number of problems. The 40-molecule burst size is much smaller than the average number of Sens molecules at the larval stage for states 3 and 4. These numbers increase to 710 and 1,760 molecules for states 3 and 4 respectively, at the larval-pupal transition. Thus, increments in bursting do not obviously translate to increments in protein number for states 3 and 4. Another problem with the burst model is the invariant ratio of Sens number observed within cells in adjacent states. It is difficult to reconcile how incremental bursting would translate to a relativistic difference in protein output.

An alternate mechanism is that the Wg and Delta signals are transduced into discrete *sens* activation states. While the states do not spatially correlate with the known concentration gradient of Wg, it is possible that the states are related to a combination of signals. If so, the distribution of the Notch ligand Delta would be an ideal candidate. Gradients of Delta have been proposed to drive sensory fate resolution in the thorax (Corson et al., 2017). Cells located in the center of thoracic stripes of proneural expression receive less Delta inhibition and therefore keep progressing to SOP fates while more lateral cells are robustly inhibited. However, there is no evidence of gradients of Delta expression in wing margin cells that resemble the thoracic stripes (Kooh et al., 1993). A more likely distribution of Delta might be that it is also discrete, since Delta is transcriptionally activated by Sens in wing margin cells (Nolo et al., 2001). Thus, cells in one Sens state would express Delta at a level that reflects a discrete state of expression different from cells in other Sens states. In turn, such Delta discretization would feedback and enhance the discretization of Sens expression. It will be interesting to determine if multimodal expression of Delta exists in wing margin cells and how their states relate to the Sens states.

Another explanation for Sens metastability might be related to the dynamics of Wg synthesis and secretion. If transcription-translation of Wg occurs in bursts, then it is possible that midline cells secrete Wg in a pulsatile manner. Nearby cells would receive and transduce the Wg signal with pulsatile dynamics, resulting in discrete time periods when transcription-translation of Sens would occur. Of course, the correct mechanism might be some combination of these various models.

Intriguingly, the metastable states have properties that are relativistic with one another. The ratio of Sens number between adjacent states is invariant across a wide range of absolute Sens protein number. And the number of Sens-positive cells in a particular state is also invariant over a wide range of Sens protein number. These relativistic properties might provide robustness against variation in absolute expression due to genetic and environmental perturbations. Indeed, in several developmental contexts it has been suggested that relative fold-changes, rather than absolute concentration changes, drive robust responses to noisy morphogen or signaling gradients, such that cells can correctly ascertain their spatial positions relative to their neighbors (Simsek and Özbudak, 2018; Lee et al., 2022). There is precedent for Wnt ligands inducing a fold-change in signal transduction to the nucleus rather than a change in absolute level (Goentoro and Kirschner, 2009). Similarly, EGF ligands have been observed to induce a constant fold-increase of 1.25 #0.11 in ERK2 nuclear levels on stimulation, despite a 4-fold variation in basal levels before stimulation (Cohen-Saidon et al., 2009). Similar features have been described for other signals, where target genes of the pathways respond to fold-changes in signal transduction (Frick et al., 2017; Lee et al., 2014) or the rate of concentration changes of key factors (Heemskerk et al., 2019). However, relativistic properties have not been observed in developmental systems exhibiting evidence of metastability. If metastability of Sens is controlled by Wg and Delta signaling, one possibility is that fold-change in signal transduction over time is translated into relativistic maintenance of the four metastable states.

Given that SOP numbers and positions are well-known to be invariant, we speculate that metastable states during fate progression aid patterning accuracy. From previous studies we know that SOP transitions are sensitive to noisy gene expression (Giri et al., 2020). The presence of metastable states suggests that cells are not allowed to freely sample all Sens levels. Instead, variation in Sens expression is restricted within a strict range, such that chance transitions to a different or SOP state are limited. Another intriguing possibility is that discrete cell states reduce the time required for developmental pattern resolution by aiding cell-competition. A single SOP is selected from a proneural field of 30-50 cells. However, neighboring cells in the procedural zone are subjected to similar levels of Wg and other pre-patterning factors. Therefore, one would expect their Sens levels to be quite similar at the start of pattern resolution. Inducing larger differences between cells by flipping some cells into higher or lower states would aid cell-competition time by requiring fewer rounds of inhibitory signaling between neighbors.

Our study raises many questions that remain unanswered. What is the nature of the metastability and what causes it? Why is it relativistic? And are the different states disposed to adopting particular terminal fates with different probabilities? These questions will be more robustly addressed once an ex vivo live imaging system of the late larval wing pouch becomes available.

## LIMITATIONS OF STUDY

The empirical results reported herein should be considered in the light of some limitations. The primary limitation being that we estimate progression of cells towards SOP fate solely by measuring their concentration of Senseless (Sens). It should be noted however, that there are other critical proneural factors such as Achaete (Ac) and Scute (Sc) and anti-neural factors such as Notch (N) and Delta (Dl) whose levels regulate the transition to SOP fate (Corson et al., 2017; Garcia-Bellido and de Celis, 2009; Hinz et al., 1994; Jafar-Nejad et al., 2006, 2003). We chose to measure Sens protein due to the abundance of literature indicating that it functions as a bistable fate switch Jafar-Nejad et al., 2003; Nolo et al., 2000; Powell et al., 2012). Further, while Ac-Sc have been shown to be important in promoting proneural competence towards SOP fate, without Sens expression, Ac-Sc are insufficient to drive selection of SOPs from proneural cells (Jafar-Nejad et al., 2003). However, in future experiments it will be important to look at the co-expression patterns of Ac-Sc as well anti-neural factors N and Dl in single cells to fully ascertain the dynamics of fate transition.

A second limitation concerns the modeling choices made to estimate the number and properties of cell states. We chose to assume four latent states underlying the observed Sens distribution data. This choice was supported by decreasing information scores up to a 4-state model. From a Bayesian perspective, if we assume that the number of latent states itself is a variable that changes from disc to disc, it is very likely that the true number of states could vary anywhere from 3 – 5 for a given disc. However, for ease of analysis and comparison we chose to fix this value at four states for both larval and larval-pupal stages. To evaluate if a parsimonious 3-state model might better fit the Sens distributions observed in larval discs, we fitted both a 3-state and 4-state model and compared the results. We observed that the additional state in the 4-state model was being generated only at the highest Sens levels. While it is possible that the 4th larval state is spurious, we opted to use a 4-state model for both stages in order to stay consistent in our analysis between larval and larval-pupal comparisons. It will be important to directly observe the latent states proposed in this study through live-imaging single cells. This would further help understand the whether cells can transition between states, and how much time a cell might spend in each of the states i.e. the metastable dynamics of cell states.

A minor limitation concerns the estimation of Sens molecule counts in single cells. We derive an empirical measure of the fluorescence to molecules conversion factor for our experiments. However, it is important to note that this is an approximation of the true conversion factor. While the relative cellular concentrations of Sens within and between discs can be accurately determined from careful image analysis, at this time we do not have a means to provide an exact estimate of the fluorescence to molecules conversion factor. Since a change in the stated conversion factor does not alter any results, this is minor limitation that does not affect the conclusions of this study.

## AUTHOR CONTRIBUTIONS

The experimental work was conceived by R.G. and R.W.C. Experimental work was performed by R.G. and S.B., under the supervision of R.W.C. D.K.P. designed, performed, and analyzed FCS experiments. Mathematical modeling was performed by R.G. All data and model analysis were performed by R.G. under the supervision of R.W.C. The manuscript was written by R.G. and R.W.C with the contribution of S.B. and D.K.P.

## ACKNOWLEDGEMENTS

Fly stocks from Hugo Bellen, Fen Biao Gao, and the Bloomington Drosophila Stock Center are gratefully appreciated. Antibodies were gifts from Hugo Bellen and purchases from the Developmental Studies Hybridoma Bank. We thank Jessica Hornick and the Biological Imaging Facility for help with imaging. Financial support was provided from the Northwestern Data Science Initiative (R.G.), Robert H. Lurie Comprehensive Cancer Center (R.G.), NIH (R35GM118144, R.W.C.), NSF (1764421, R.W.C.), and the Simons Foundation (597491, R.W.C.).

## STAR METHODS

### RESOURCE AVAILABILITY

#### Lead contact

Further information and requests for resources and reagents should be directed to and will be fulfilled by the lead contact, Richard Carthew.

#### Materials availability

This study did not generate new unique reagents.

#### Data and code availability

- All data reported in this paper will be shared by the lead contact upon request.
- All original code has been deposited on Github and is publicly available as of the date of publication. The DOI is listed in the Key Resources Table.
- Any additional information required to reanalyze the data reported in this paper is available from the lead contact upon request.

### EXPERIMENTAL MODEL AND SUBJECT DETAILS

For all experiments, *Drosophila melanogaster* was raised using standard lab conditions and food. Stocks were either obtained from the Bloomington Stock Center, from listed labs, or were derived in our laboratory (RWC). Wild-type refers to stock *w^1118^*. MicroRNA miR-9a null mutant stocks *mir9a^E39^* and *mir9a^J22^* were provided courtesy of the Gao Lab (Li et al., 2006). The *sens* mutant alleles used *sens^E1^* and *sens^E2^* are protein null mutants (Nolo et al., 2000, 2001). A list of all mutants and transgenics used in this study is in the Key Resources Table. All experiments used female animals unless stated otherwise. Animals were staged based on morphology. The larval-pupal transition stage is one marked by the animals appearing as white prepupae. This stage lasts for only 45-60 minutes. The late larval stage is one marked by the animals as large wandering larvae with clear guts. This stage lasts for about 1-2 hours.

N-terminal 3xFlag-TEV-StrepII-sfGFP-FlAsH tagged *sens,* originally generated from the CH322-01N16 BAC, was a kind gift from K. Venken and H. Bellen (Venken et al., 2006, 2009). It has been shown to rescue *sens^E1^* and *sens^E2^* mutations (Cassidy et al., 2013; Venken et al., 2006). Generation of the 3xFlag-TEV-StrepII-mCherry-FlAsH tagged *sens,* as well as *miR-9a* binding site mutant alleles of the tagged *sens* transgenes has been described previously (Giri et al., 2020). Cloning details are available on request. All BACs were integrated at PBacy+-attP-3BVK00037 (22A3) landing site by phiC31 recombination (Venken et al., 2006). Transgenic lines were crossed with *sens* mutant lines to construct stocks in which *sens* transgenes were present in a *sens^E1^*/ *sens^E2^* trans-heterozygous mutant background. Thus, the only Sens protein expressed from these animals came from the transgenes.

### METHOD DETAILS

#### Fluorescence microscopy and quantification

Wing discs from staged animals were dissected out in ice-cold Phosphate Buffered Saline (PBS). Discs were fixed in 4% (w/v) paraformaldehyde in PBS for 20 minutes at 25°C and washed with PBS containing 0.3% (v/v) Tween-20. Then they were stained with 0.5 μg/ml 4′,6-diamidino-2-phenylindole (DAPI, Thermo Scientific), and mounted in Vectashield (Vector Labs). Discs were mounted apical-side up and imaged with identical settings using a Leica TCS SP5 inverted confocal microscope. Images were acquired with a 100x (NA – 1.44) oil-immersion objective, at 2048 x 2048 resolution, with a 75 nm x-y pixel size, and 0.42 μm z separation. Scans were collected bidirectionally at 400 MHz, and 6x line in the same session, for consistency in data quality. Image segmentation, quantification, and conversion to absolute numbers of Sens molecules was carried out using a custom computational pipeline as described (Giri et al., 2020). Briefly, for each wing disc five optical slices containing proneural cells were chosen for imaging and analysis. Cells nuclei were computationally segmented in each slice using a custom MATLAB script (Peláez et al., 2015; Qi et al., 2013). For each segmented nucleus, the relative position of its centroid in x and y was calculated, along with the average sfGFP signal intensity in the nuclear area. The majority of cells imaged did not fall within the proneural region and therefore displayed background levels of fluorescence scattered around some mean level with Sens expressing cells present in the right-hand tail of the distribution. Since the background signal varied slightly from disc to disc, we calculated a disc-specific “mean channel background”. This was done by fitting a Gaussian distribution to the population and finding the mean of that fit. In order to separate Sens positive cells, we chose a cut-off percentile based on the Gaussian distribution, below which cells were deemed Sens negative. We set this cut-off at the 99^th^ percentile for all analysis, at both the larval and larval-pupal stage. In order to ensure that outliers did not skew the estimate of this mean background, the full distribution was first trimmed before calculating the parameters of the background Gaussian fit. Finally, the fluourescence of each cell in the disc was normalized by subtracting out the value of the 99^th^ percentile background fluourescence calculated for that disc. The background cutoff was not significantly different between the larval and larval-pupal stages as determined by a t-test (p=0.23). This method yielded the expected pattern of Sens-positive cells distributed in two stripes across the dorsoventral midline of the wing disc. On average, 1,000 ± 200 Sens-positive segmented nuclear-slices were quantified from each imaged wing disc, and a total of 6 to 12 discs were imaged for each condition. On average a single cell was present in 4 nuclear-slices, corresponding to 250 # 50 Sens-positive cells per disc.

#### Immunohistochemistry

Discs were dissected, and fixed as above, before incubating with the primary antibodies of interest. Tissues were washed thrice for 5-10 minutes each and incubated with the appropriate fluorescent secondary antibodies (diluted 1:250) for 1 hour. They were then stained with DAPI, washed in PBS-Tween, and mounted for imaging. Guinea pig anti-Sens antibody (gift from H. Bellen) was used at 1:1000 dilution. Mouse monoclonal anti-Wg 4D4 was obtained from the Developmental Studies Hybridoma Bank (DSHB) and used at a 1:25 dilution. Secondary antibody Alexa Fluor 488 conjugated Goat anti-guinea pig IgG was used at 1:250 dilution, and Alexa Fluor 546 conjugated Goat anti-mouse IgG was used at 1:200 dilution.

#### FCS sample calibration and measurement*s*

A previously determined FCS calibration factor was used to convert confocal-measured nuclear tagged Sens fluorescence into nanomolar concentrations (Giri et al., 2020). Subsequently concentrations were converted into numbers of Sens molecules by multiplying with average cell volumes. The protocol for FCS is reproduced here for reference.

White pre-pupal wing discs of genotype *sfGFP-sens, sens^E1^/mCherry-sens, sens^E2^* or *sfGFP-sens, sens^E2^/mCherry-sens, sens^E1^* were dissected in PBS and sunken into LabTek 8-well chambered slides containing 400 l PBS per well (Papadopoulos et al., 2019). Discs were positioned such that the pouch region was facing the bottom of the well to be imaged. FCS measurements were made using an inverted Zeiss LSM780, Confocor 3 instrument with APD detectors. A water immersion 40x objective with numerical aperture of 1.2 (which is optimal for FCS measurements) was used throughout. Fast image scanning was utilized for identification of cell nuclei to be measured by FCS. Prior to each session, we used 10 nM dilute solutions of Alexa488 and CF586 dyes to calculate the average number of particles, the diffusion time and define the structural parameters. Using these we calibrated the Observation Volume Element (OVE) whose volume can approximated by a prolate ellipsoid . Measurements were performed in Sensory Organ Precursor cells (SOPs), as well as first and second order neighbors, residing dorsally or ventrally of the SOP cell. Measurements were subjected to analysis and fitting, using a two components model for three-dimensional diffusion and triplet correction as follows:

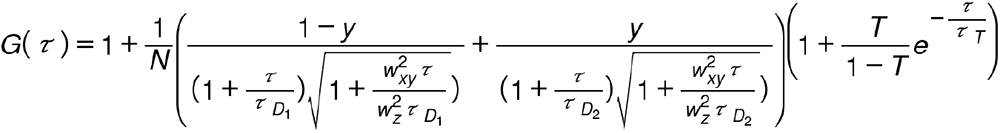

In the above equation, *N* is the average number of molecules in the OVE; *y* is the fraction of the slowly moving sfGFP-Sens or mCherry-Sens molecules; 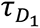 is the diffusion time of the free sfGFP-Sens or mCherry-Sens molecules; 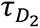 is the diffusion time of sfGFP-Sens or mCherry-Sens molecules undergoing nonspecific interactions with chromatin; *w_xy_* and *w_z_* are radial and axial parameters, respectively, related to spatial properties of the OVE; *T* is the average equilibrium fraction of molecules in the triplet state; and τ_T_ the triplet correlation time related to rate constants for intersystem crossing and the triplet decay. Spatial properties of the detection volume, represented by the square of the ratio of the axial and radial parameters 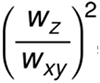, are determined in calibration measurements performed using a solution of A488 and CF568 for which the diffusion coefficient (*D*) is known. The diffusion time, τ_*D*_, measured by FCS, is related to the translation diffusion coefficient D by 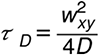. Measurements which exhibited marked bleaching, low counts per molecule (CPM below 0.5 kHz per molecule per second) or measurements in which fitting of the auto-correlation curves (ACCs) yielded abnormally high or low diffusion times of the fast FCS component, were excluded from the analysis.

From FCS measurements, the highest Sens expression from either allele (sfGFP-Sens or mCherry-Sens) was no greater than 250 nM, corresponding to approximately 25 RFU obtained from imaging. Additionally, the majority of first or second order neighbors of SOP cells were observed to lie below an allelic expression level of 125 nM, again corresponding to approximately 12.5 RFU in images.

This provided an approximate conversion factor of 1 RFU equivalent to 10 nM. To validate this estimate, we classified Sens-positive cells as SOPs (total Sens > 250 nM), proximal neighbors (Sens between 50-250nM), or distant neighbors (Sens < 50 nM) and mapped them to raw images. An accurate estimate of a conversion factor would to recreate the expected expression pattern of Sens, i.e. SOP-fated cells dispersed periodically along both sides of the wing margin surrounded by first and second order neighbours. We reproducibly observed this pattern for a conversion factor of 10 nM/RFU, but not when it was varied three-fold higher or lower. Since the average wing disc nuclear volume is 23 x 10^-15^ L (Papadopoulos et al., 2019), 1 RFU is estimated to correspond to ∼138 Sens molecules. We refer the reader to Figure 1G and Figure 1 – figure supplement 3 from Giri et al., 2020 for further details.

To ascertain that the fluorescence to molecules conversion factor estimated from *sfGFP-sens/mCherry-sens* animals could be faithfully extrapolated to homozygous *sfGFP-sens* animals, we verified that there were no FRET interactions between sfGFP-Sens and mCherry-Sens *in vivo.* This was done by tuning the 488nm laser to excite sfGFP-Sens incrementally and measuring both green and red photon emission from the cell (Fig. S2A). We also ascertained the lack of FRET or other interfering processes by gradually expanding the FCS observation volume and verifying the expected linear increases in molecule counts and diffusion times. Finally, we confirmed that tagged Sens species diffusion rates do not change with increases in Sens nuclear concentration (Fig. S2B,C) or with developmental stage (Fig. S2D).

#### Cell distance estimation

Cell distances from the wing pouch dorsoventral midline were estimated by approximating a curved line representing the midline for each disc, and subsequently calculating the shortest straight-line distance between this approximated margin and the cell centroid as obtained from the automated image-quantification pipeline.

#### Temperature perturbation

Animals were grown in either 18°C or 29°C incubators until the desired stage of development was reached, at which point the wing discs were rapidly dissected, fixed and processed as described above.

#### Mixture model analysis and probability density functions

Probability density functions were calculated using a kernel density estimation. The appropriate value for the bandwidth parameter was estimated using either Scott’s rule or Silverman’s rule. Both estimators were independently used and both gave highly similar results (Fig. S3). For mixture model analysis, the R package mixtools and Python package sklearn were used. Single-cell Sens distributions in individual discs were assumed to be composed of *k* Guassian sub-distributions. The best-fit value of *k* was determined by testing mixture models with different values of *k* and comparing the change in relative information score. Both the Akaike Information Criterion (AIC) and Bayesian Information Criterion (BIC) were evaluated since the AIC score is known to be biased towards over-parameterization. The least parametrized model that yielded sufficiently low AIC and BIC scores across several discs from both larval and larval to pupal transition stages was determined empirically to be 4. Using parameter *k* = 4, the mean, standard deviation and proportion were determined for each sub-population within a population of Sens-positive cells obtained from each individual wing pouch. The values of mean, standard deviation, and proportion were averaged from six to twelve wing pouches per treatment (Table 1). We observed that each subpopulation mean was separated from the adjacent subpopulation’s mean by a value greater than the sum of their standard deviations (Table 1). Moreover, each sub-population’s coefficient of variation was fairly similar to the other sub-populations’, regardless of sample or stage (Table 1). The number of cells in each state was calculated by assuming that each cell appears in approximately 4 out of 5 optical slices imaged per pouch and multiplying the total number of Sens-positive cells in the pouch by the proportion value of each state for that pouch.

#### Mathematical model for patterning

The mathematical model for self-organization of *Drosophila* proneural tissue and fate selection is as described by Corson et. al. (2017) as

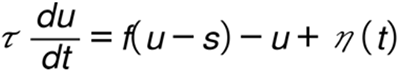

The model variable *u* describes the fate potential of each cell, which is assumed analogous to the concentration of proneural proteins such as Sens. Variable *s* describes the cumulative level of lateral inhibition signal received by the cell. Thus, *s* can be thought to be analogous to Delta-activated Notch inhibition of Sens and other proneural factors. Although *u* is not formally the level of an individual proneural factor like Sens, the model allows for the variable to approximate it. To simulate patterning of the wing margin, we prescribed the system as a single row of cells in which the *u* evolves over time *t*, and resolves into an epidermal state (*u* = 0) or SOP state (*u* = 1). Variable *τ* sets the the time-scale of cell dynamics.

The functional form and parameter values describing the function *f(u−s)* which integrates the activating and inhibiting signals, were kept identical to Corson et al (2017). The model was simplified to a 1-dimensional row of 20 cells with periodic boundary conditions, and 100 simulations were run in parallel for each condition. The Brownian noise term *η(t)* was set at 1% relative to *u* (process noise = 0.01) for the simulation results presented in Figure 2. This term describes the level of stochasticity in the dynamics of cell state *u* encoded in the model. A total of 250 time-steps were simulated in order to ensure convergence, however cell-state resolution was generally observed within 25-50 time steps, therefore Figure 2 shows the time-evolving distribution at every third time-step in the series from steps 1-30.

As shown, for all cells, *u* first reaches an intermediate steady state that lasts for some variable amount of time until *u* bifurcates towards the 0 or 1 final states. This transient steady state represents an unstable one, since the system has a saddle-node bifurcation (Fig. 2A). In model simulations, the time course of cell state shows a gradual progression of cells toward the SOP state, accompanied by a continuous branching off of cells that relax to zero *u* (Fig. 2B). The final pattern of cell states resembles that observed for the wing margin SOPs; a row of periodically spaced SOPs (Fig. 2C). Code and all parameter values for explicitly generating the simulations and figure plots is available at https://github.com/ritika-giri/metastable-states-Sens.

### QUANTIFICATION AND STATISTICAL ANALYSIS

Each experimental condition was assayed by analyzing between 6-12 individual wing pouches. Within each wing pouch we identified approximately 1,000 segmented cells that were deemed ‘Sens-positive’. For histogram comparison between conditions, all segmented Sens-positive cells for a given condition were pooled and 5,000 cells were randomly sampled to generate probability distributions. For mixture analysis, all Sens-positive cells of a pouch were used to estimate subpopulation parameters. Estimates for individual pouches are shown directly in the relevant box plot graphs. To estimate the number of real cells, the segmented cell count was divided by 4 with the assumption that each cell is present in roughly 4 out 5 optical slices assayed for each wing pouch. Wilcoxon’s test was used to calculate a p-value for all comparisons, unless stated otherwise. To calculate a p-value for the ratios in mean Sens number between states, a Z test was administered. For all experiments, there was no exclusion of any data or subjects.

## KEY RESOURCES TABLE

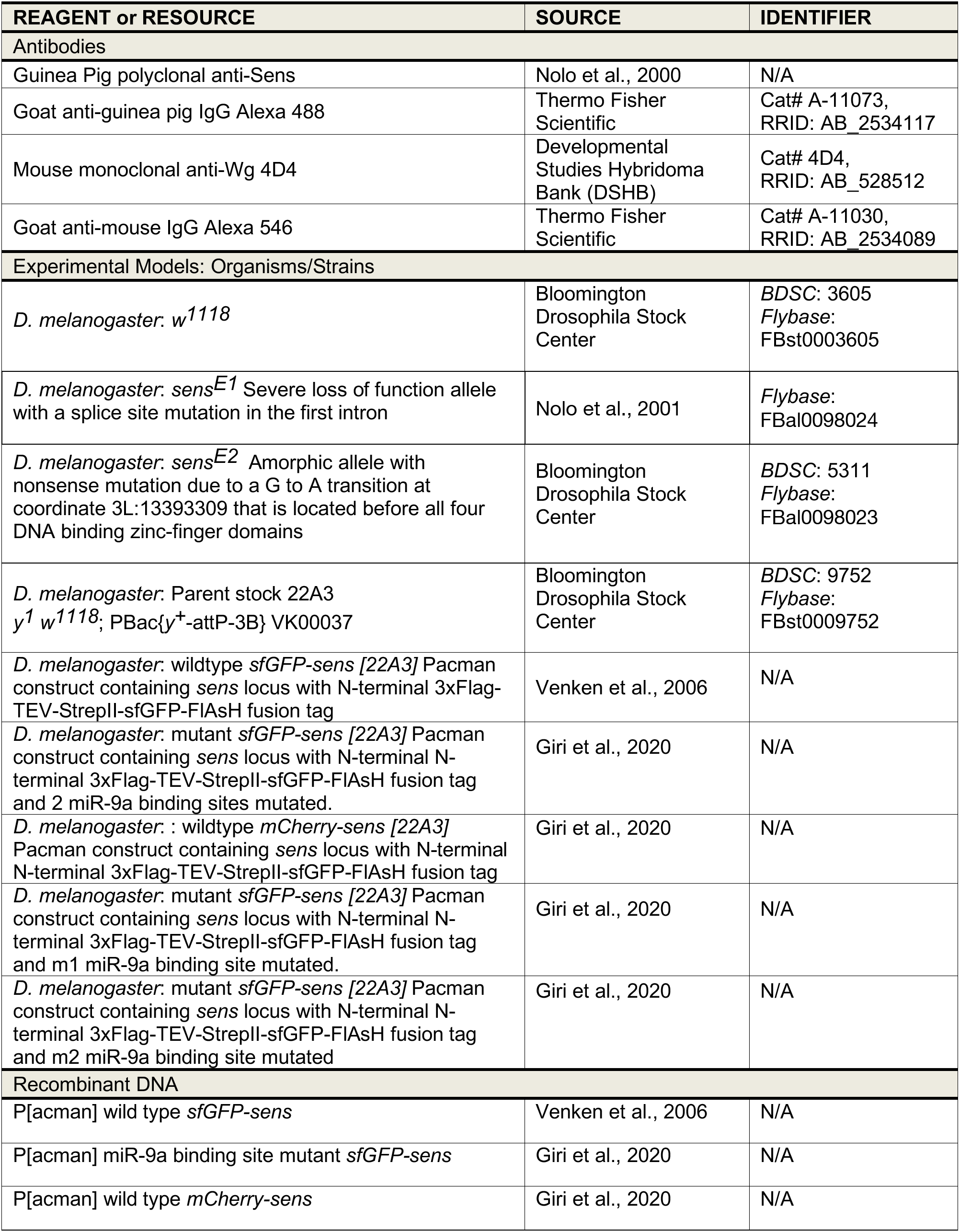
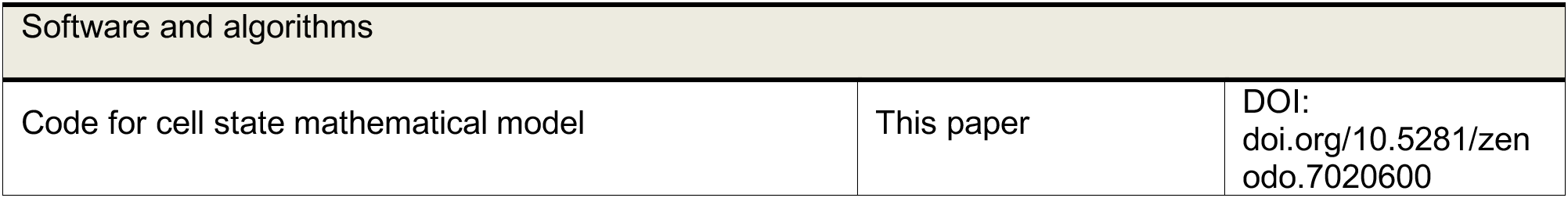

| REAGENT or RESOURCE | SOURCE | IDENTIFIER |
| --- | --- | --- |
| Antibodies |  |  |
| Guinea Pig polyclonal anti-Sens | Nolo et al., 2000 | N/A |
| Goat anti-guinea pig IgG Alexa 488 | Thermo Fisher Scientific | Cat# A-11073, RRID: AB_2534117 |
| Mouse monoclonal anti-Wg 4D4 | Developmental Studies Hybridoma Bank (DSHB) | Cat# 4D4, RRID: AB_528512 |
| Goat anti-mouse IgG Alexa 546 | Thermo Fisher Scientific | Cat# A-11030, RRID: AB_2534089 |
| Experimental Models: Organisms/Strains |  |  |
| *D. melanogaster*: *w*<sup>1118</sup> | Bloomington Drosophila Stock Center | BDSC: 3605<br>Flybase: FBst0003605 |
| *D. melanogaster*: *sens*<sup>E1</sup> Severe loss of function allele with a splice site mutation in the first intron | Nolo et al., 2001 | Flybase: FBal0098024 |
| *D. melanogaster*: *sens*<sup>E2</sup> Amorphic allele with nonsense mutation due to a G to A transition at coordinate 3L:13393309 that is located before all four DNA binding zinc-finger domains | Bloomington Drosophila Stock Center | BDSC: 5311<br>Flybase: FBal0098023 |
| *D. melanogaster*: Parent stock 22A3 *y*<sup>1</sup> *w*<sup>1118</sup>; PBac{*y*<sup>+</sup>-attP-3B} VK00037 | Bloomington Drosophila Stock Center | BDSC: 9752<br>Flybase: FBst0009752 |
| *D. melanogaster*: wildtype *sfGFP-sens* [22A3] Pacman construct containing *sens* locus with N-terminal 3xFlag-TEV-StrepII-sfGFP-FIAsH fusion tag | Venken et al., 2006 | N/A |
| *D. melanogaster*: mutant *sfGFP-sens* [22A3] Pacman construct containing *sens* locus with N-terminal N-terminal 3xFlag-TEV-StrepII-sfGFP-FIAsH fusion tag and 2 miR-9a binding sites mutated. | Giri et al., 2020 | N/A |
| *D. melanogaster*: : wildtype *mCherry-sens* [22A3] Pacman construct containing *sens* locus with N-terminal N-terminal 3xFlag-TEV-StrepII-sfGFP-FIAsH fusion tag | Giri et al., 2020 | N/A |
| *D. melanogaster*: mutant *sfGFP-sens* [22A3] Pacman construct containing *sens* locus with N-terminal N-terminal 3xFlag-TEV-StrepII-sfGFP-FIAsH fusion tag and m1 miR-9a binding site mutated. | Giri et al., 2020 | N/A |
| *D. melanogaster*: mutant *sfGFP-sens* [22A3] Pacman construct containing *sens* locus with N-terminal N-terminal 3xFlag-TEV-StrepII-sfGFP-FIAsH fusion tag and m2 miR-9a binding site mutated | Giri et al., 2020 | N/A |
| Recombinant DNA |  |  |
| P[acman] wild type *sfGFP-sens* | Venken et al., 2006 | N/A |
| P[acman] miR-9a binding site mutant *sfGFP-sens* | Giri et al., 2020 | N/A |
| P[acman] wild type *mCherry-sens* | Giri et al., 2020 | N/A |
| Software and algorithms |  |  |
| Code for cell state mathematical model | This paper | DOI:<br>doi.org/10.5281/zenodo.7020600 |

---

**Figure S1.**
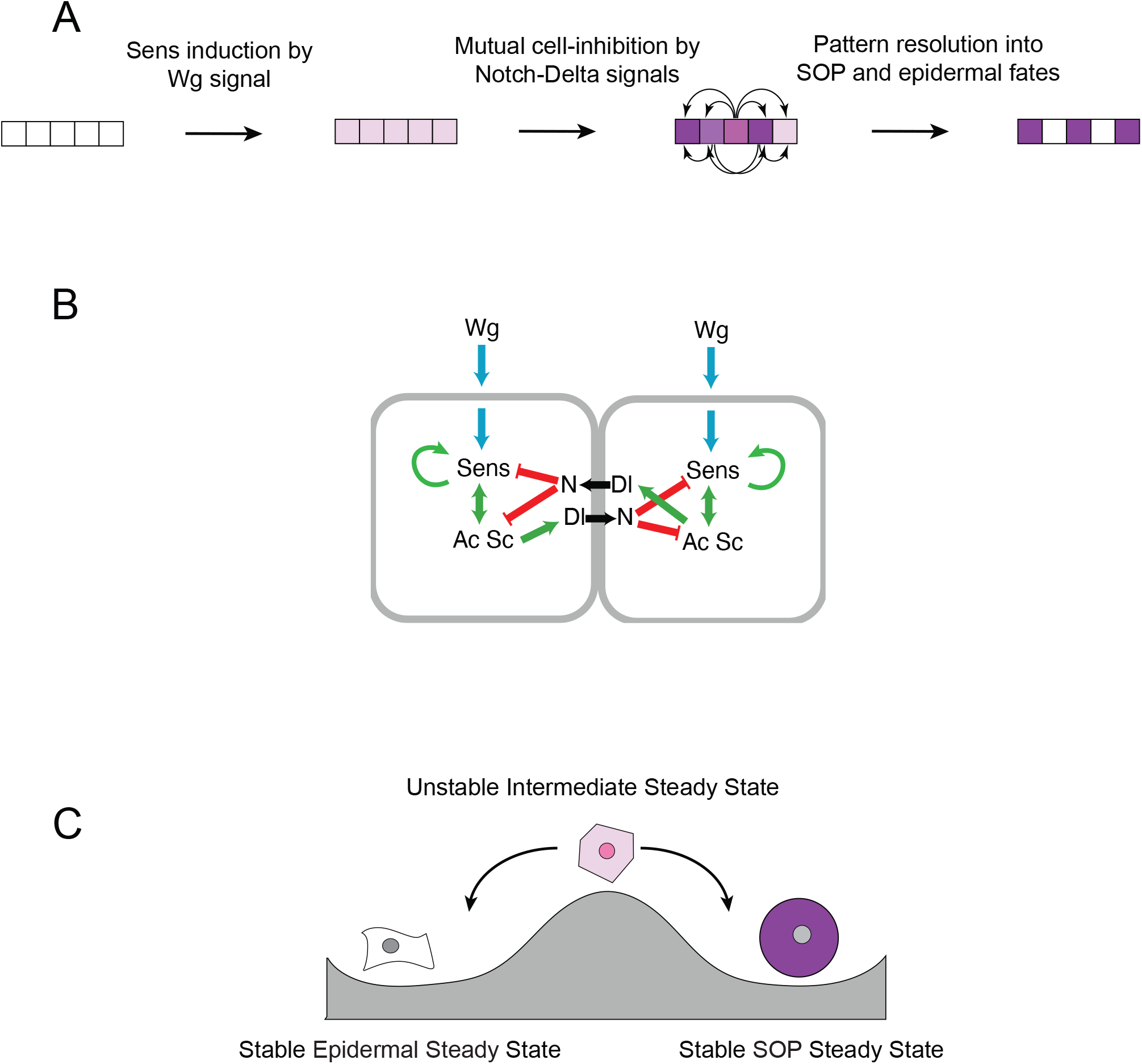
Sens expression dynamics, related to. **Figure 1. (A)** Steps in the process of resolving cell fate decisions with Sens. **(B)** Schematic of the gene regulatory network that operates in proneural cells at the anterior wing margin. **(C)** Cells undergoing the sensory versus epidermal fate decision are thought to exhibit bistability. An intermediate level of Sens expression is an unstable cell state can resolve into one of two stable states: a high Sens expressing SOP state, or a low Sens expressing epidermal state.

**Figure S2.**
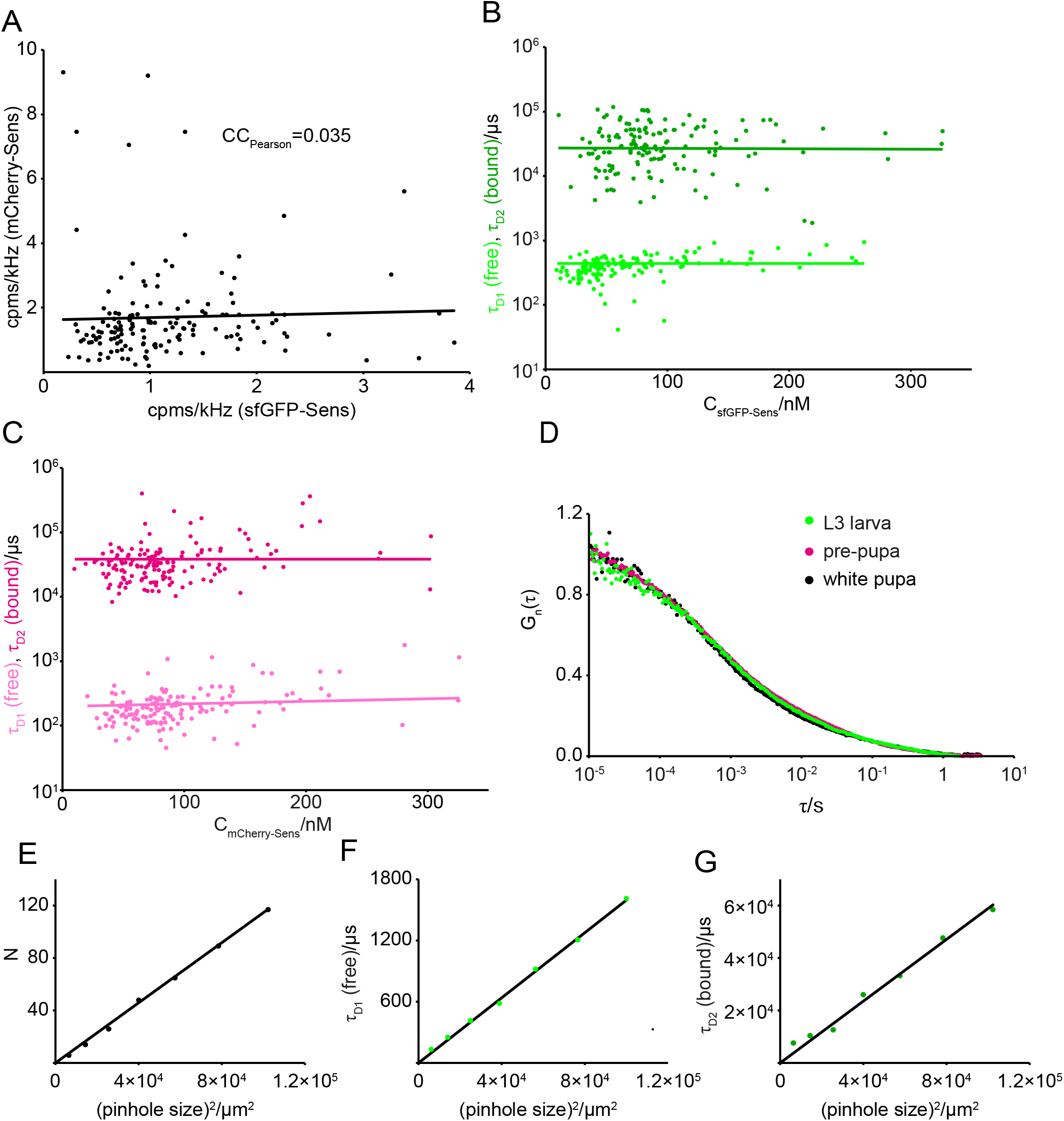
Fluorescence correlation spectroscopy analysis, related to Figure 3. **(A)** Correlation between sfGFP-Sens and mCherry-Sens photon counts per molecules and second (cpms) in transheterozygous *sfGFP-sens/mCherry-sens* wing imaginal discs in vivo. High sfGFP-Sens cpms is not accompanied by high mCherry-Sens cpms, indicating that no appreciable FRET from sfGFP-Sens to mCherry-Sens molecules takes place in vivo. **(B,C)** Characteristic decay times *τ_1_* and *τ_2_* representing freely-dif-fusing and bound sfGFP-Sens **(B)** and mCherry-Sens **(C)** molecules in vivo as a function of the total concentration of Sens in individual cells. No changes in neither sfGFP-Sens nor mCherry-Sens diffusion times were observed upon changes in nuclear concentration. **(D)** Autocorrelation curves normalized at the zero-th lag time display overlapping decay times indicating similar diffusion and fractions of chromatin-bound and freely diffusing sfGFP-Sens molecules across late-larval and early-pupal developmental stages. Shown here are average curves for FCS measurements of sfGFP-Sens from larval (green), larval-pupal (magenta) and early pupal (black) wing discs. **(E–G)** Pinhole experiments to assess effect of the enlargement of the detection volume on the characteristic decay times *τ_1_*and *τ_2_*. Pinhole size was incrementally increased to expand the FCS detection volume and the number of molecules N **(E)**, *τ_1_* **(F)**, and *τ_2_* **(G)** derived by fitting are plotted as a function of the squared pinhole size in a transheterozygous *sfGFP-sens/mCherry-sens* animal. All three parameters increase linearly with increasing detection volumes, indicating that molecular movement is the predominant cause underlying the fluorescence intensity fluctuations recorded by FCS.

**Figure S3.**
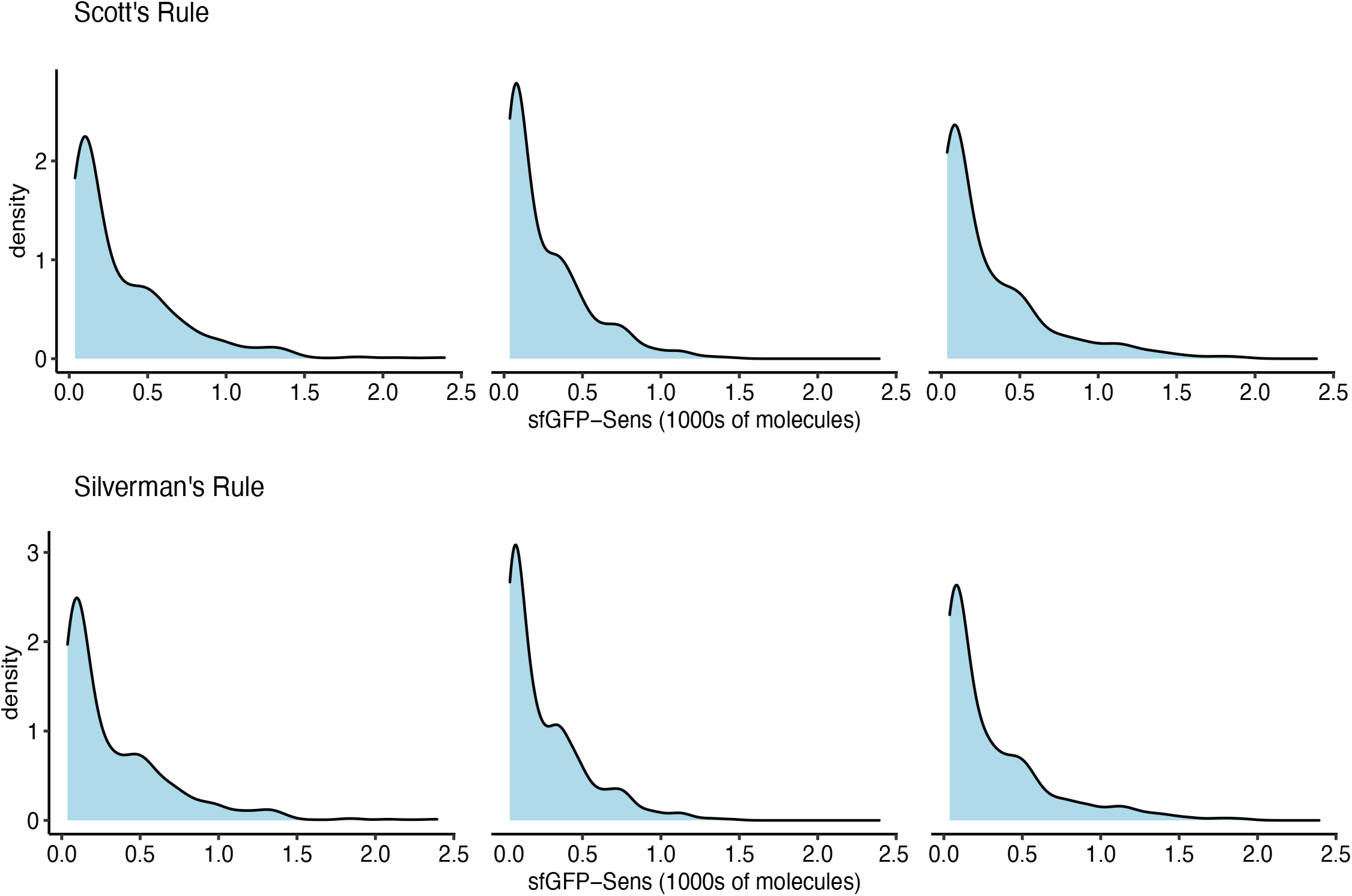
Kernel density estimations of three populations of Sens-positive cells, related to Figure 3. From left to right, each chart displays a population of Sens-positive cells from an individual wing pouch. To show that the probability density distribution qualitatively appears multimodal regardless of bandwidth parameter selection, the parameter was chosen either using Scott’s (top) or Silverman’s (bottom) rule.

**Figure S4.**
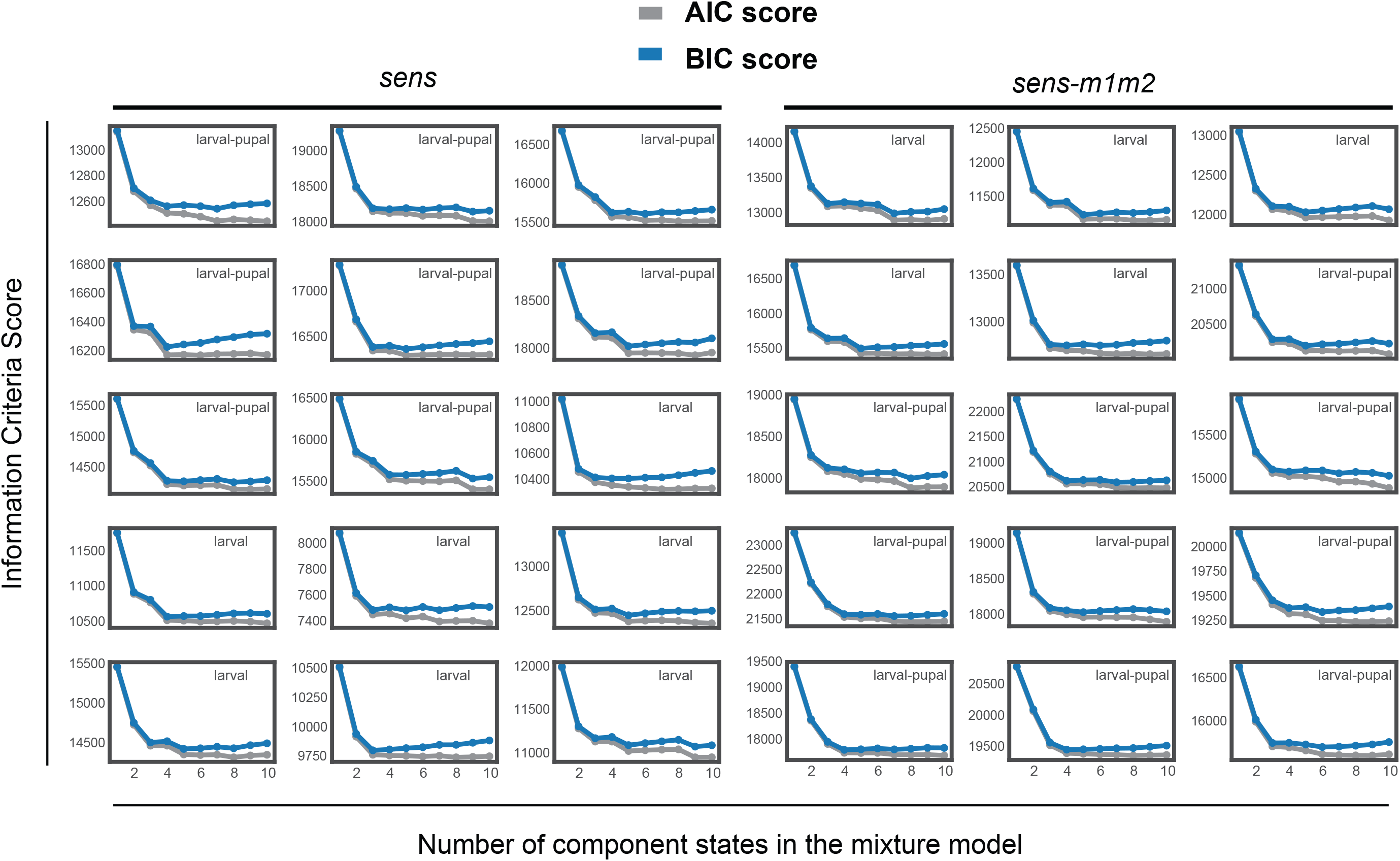
AIC and BIC scores for Gaussian mixture models with different numbers of states in the model, related to Figure 3. Sens data from individual wing pouches were independently assessed using the AIC and BIC scores. These tested the relative quality of statistical models built with different values of the parameter *k*, which defines the number of states within a parent distribution. Models with a *k* value that generates the lowest AIC and BIC score are deemed to be best-fit, in that they are best at describing the data without overfitting by having higher values of *k*. Shown in each panel are scores for individual wing pouch samples used in experiments described in Figure 3 and Figure 5A-D for wild type *sens* (left) and miR-9a binding site mutant *sens-m1m2* (right) genotypes, respectively.

**Figure S5.**
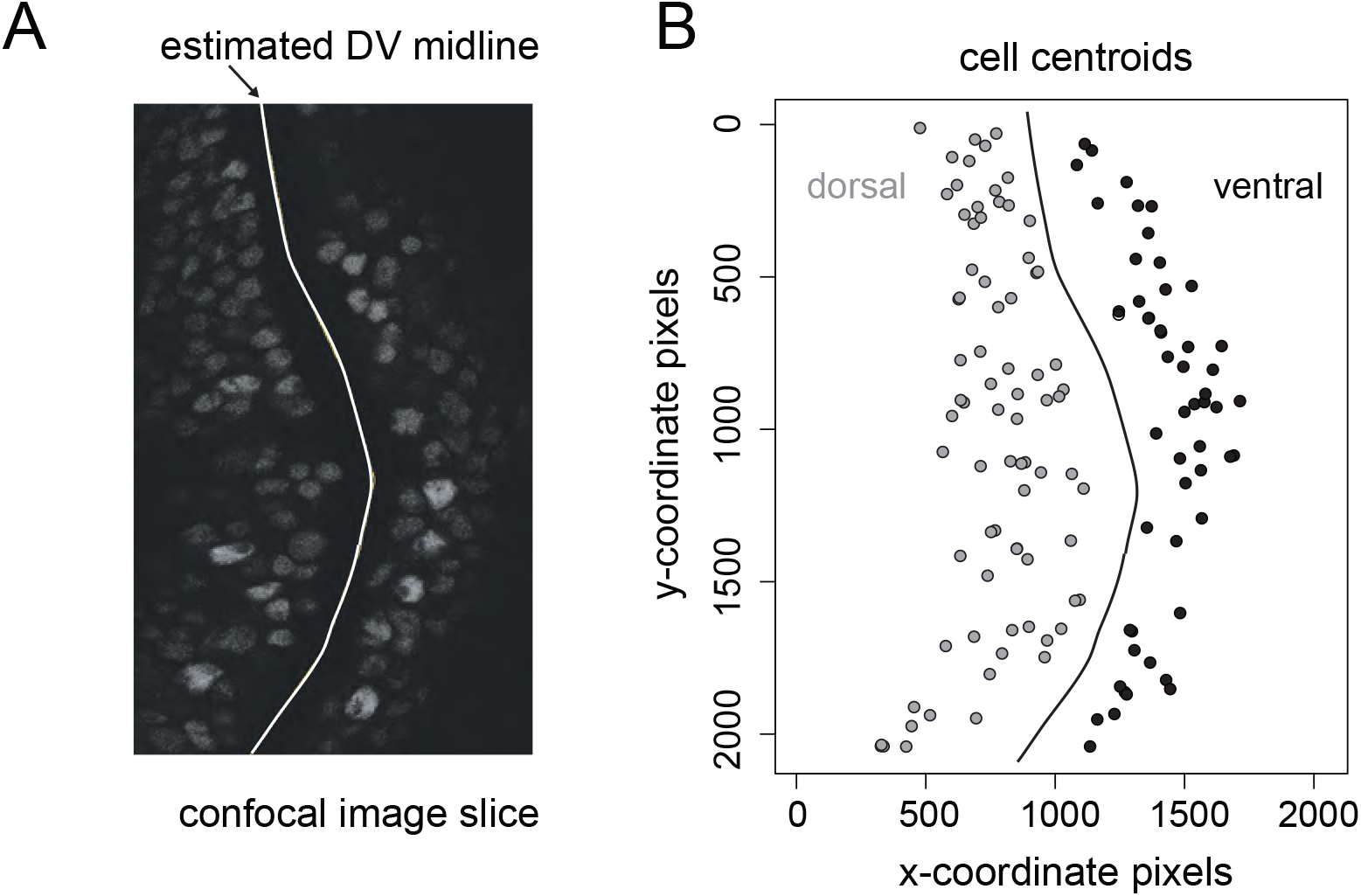
Estimation of the shortest distance from a cell to the dorsoventral midline, related to Figure 4. **(A)** A fluorescence image slice of a wing pouch expressing sfGFP-Sens is shown. To calculate cell distance from the midline, pixel coordinates making up the dorsoventral midline were estimated manually and delineated in white. **(B)** Cell-centroids of each segmented cell were mapped. Dorsal compartment cells are marked in gray, ventral in black for reference. Cell distance from the midline was calculated as the shortest Euclidean distance from the cell centroid to the estimated midline and converted to micrometers based on the imaging scale factor.

**Figure S6.**
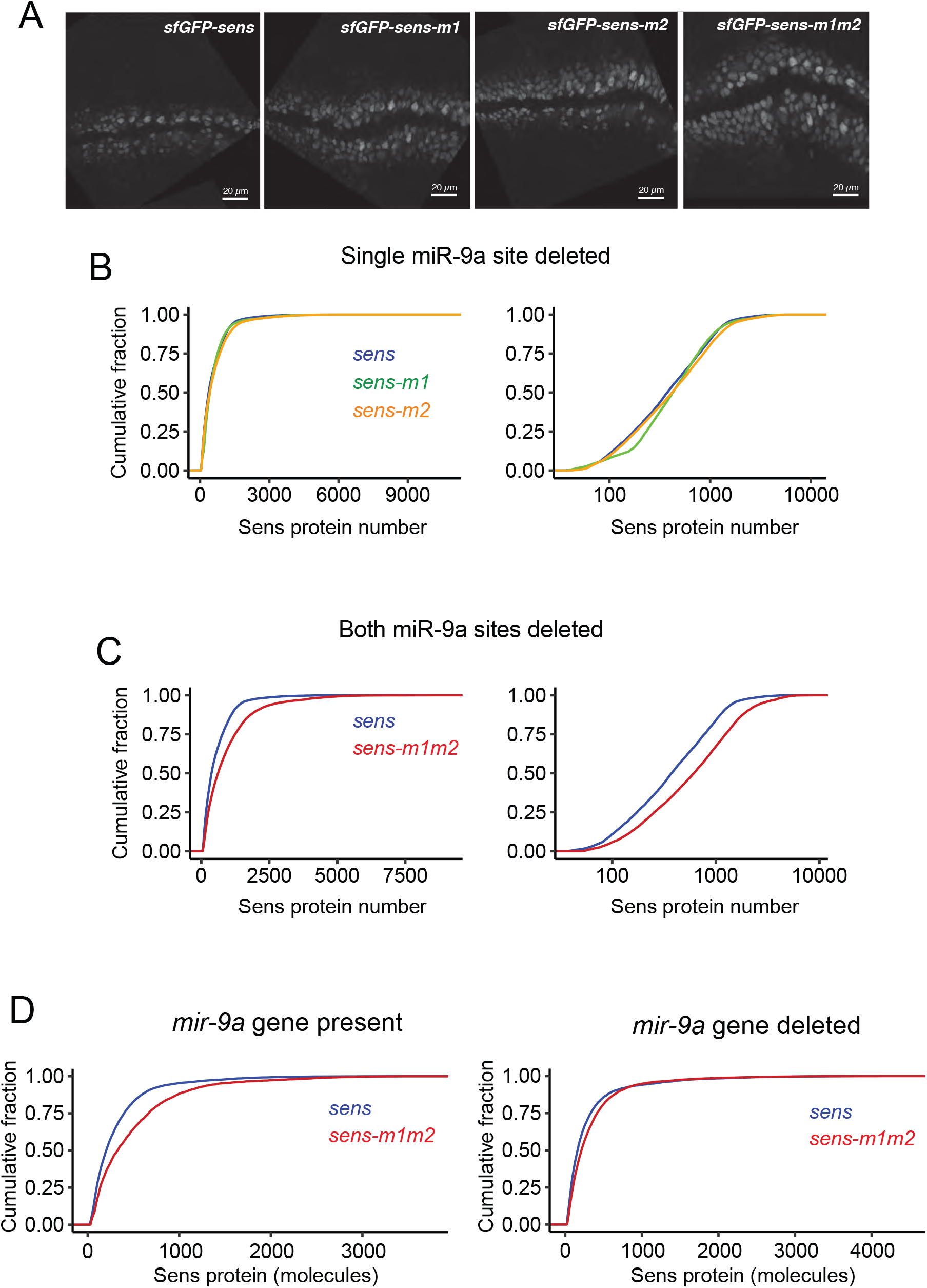
Mutation of miR-9a binding sites in sens increases Sens protein output, related to Figure 5. **(A)** Sens expression patterns for different miR-9a binding-site mutant *sens* alleles. Fluorescence from sfGFP-Sens in larval-pupal wing pouches is shown. Sens expression was assayed for different combinations of miR-9a binding-site mutations in the *sens* 3’UTR in an endogenous *sens* null background. All four alleles rescue *sens* expression. Wild-type like transgene *sfGFP-sens* had both miR-9a sites intact, site 1 and site 2 were deleted in *sfGFP-sens-m1* and *sfGFP-sens-m2* alleles respectively. Both sites were deleted in allele *sfGFP-sens-m1m2*. **(B,C)** Cumulative density plots show the distributions of Sens protein number for cells expressing various *sens* alleles. Linear scale for Sens number is on the left, log scale is on the right. **(B)** Single-site mutants *sens-m1* and *sens-m2* are compared to wild-type *sens*. **(C)** Sens protein output increases when both miR-9a binding sites are deleted in *sens-m1m2*, relative to wild-type *sens*. **(D)** Sens protein output from the wild-type *sens* and mutant *sens-m1m2* alleles was measured in the presence (left) or absence (right) of the endogenous *mir-9a* gene. The cumulative density plot shows the difference in protein output between wild-type and mutant transgenes. When mir-9a is present, wild-type Sens expression is lower compared to the m1m2 mutant. However, this difference is greatly diminished when endogenous miR-9a is not present.

**Figure S7.**
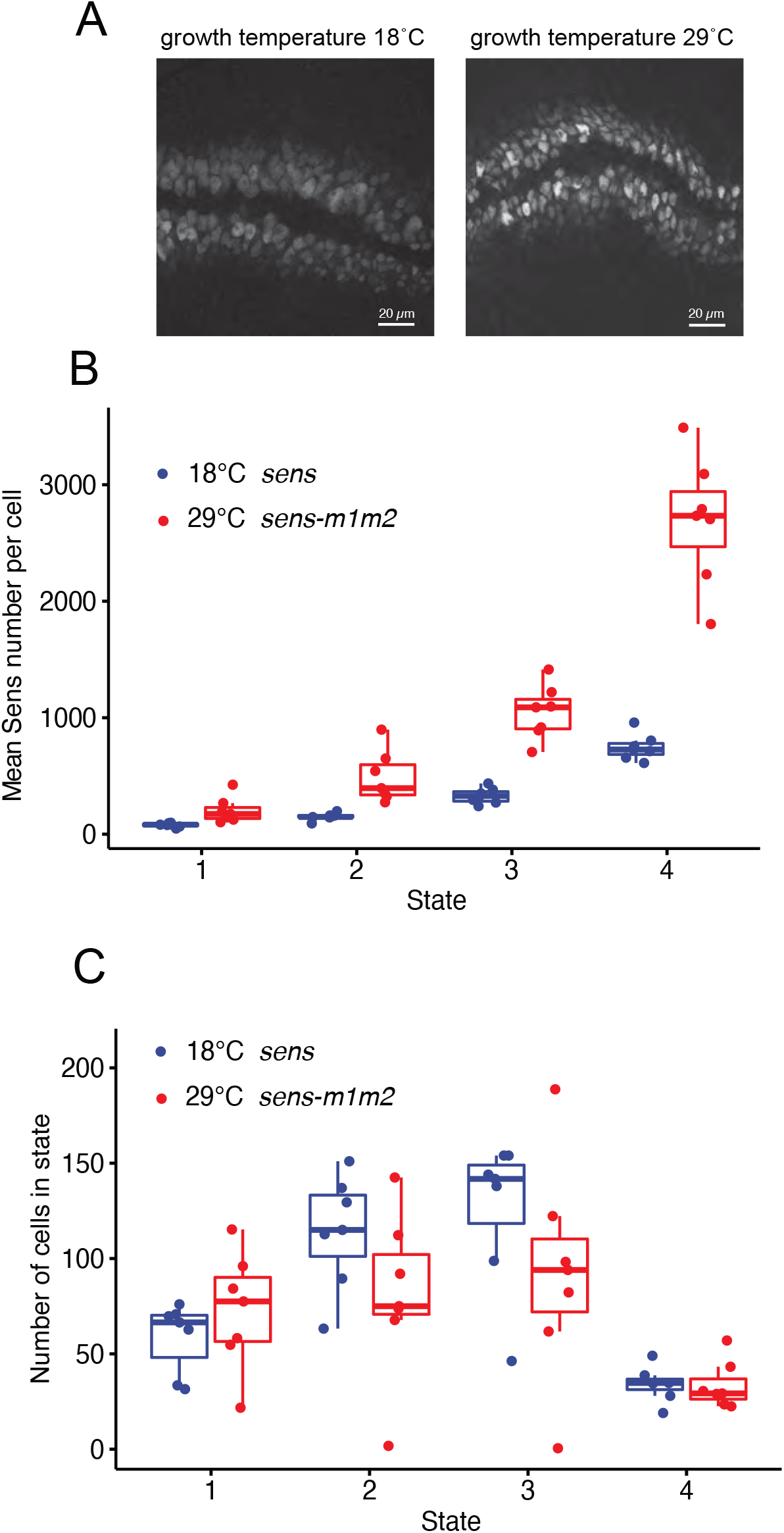
Combined change in miR-9a regulation and growth temperature have little effect on cell-state properties, related to Figure 5. **(A)** Representative larval-pupal wing pouch images showing sfGFP-Sens expression from animals raised at either 18°C or 29°C. **(B)** Sens protein number for cells in the four states. Each dot represents a mean estimate of protein number derived from one larval-pupal stage wing pouch. Multiple samples were analyzed for each condition. Wing pouches are from wildtype *sens* animals grown at 18°C (blue) or *sens-m1m2* mutant animals grown at 29°C (red). Boxplots show the median and interquartile range (25 – 75%), and whiskers represent the 1.5x interquartile range. The differences in Sens number between *sens* at 18°C and *sens-m1m2* at 29°C for each state were all significant (*p* < 0.001, Wilcoxon test). **(C)** Estimated number of cells within each state for the four states. Each dot represents an estimate derived from one larval-pupal stage wing pouch. Wing pouches are from wildtype *sens* animals grown at 18°C (blue) or *sens-m1m2* mutant animals grown at 29°C (red). Boxplots show the median and interquartile range (25 – 75%), and whiskers represent the 1.5x interquartile range. The differences in cell number between *sens* at 18°C and *sens-m1m2* at 29°C for each state were not significant (*p* > 0.15, Wilcoxon test).

**Figure S8.**
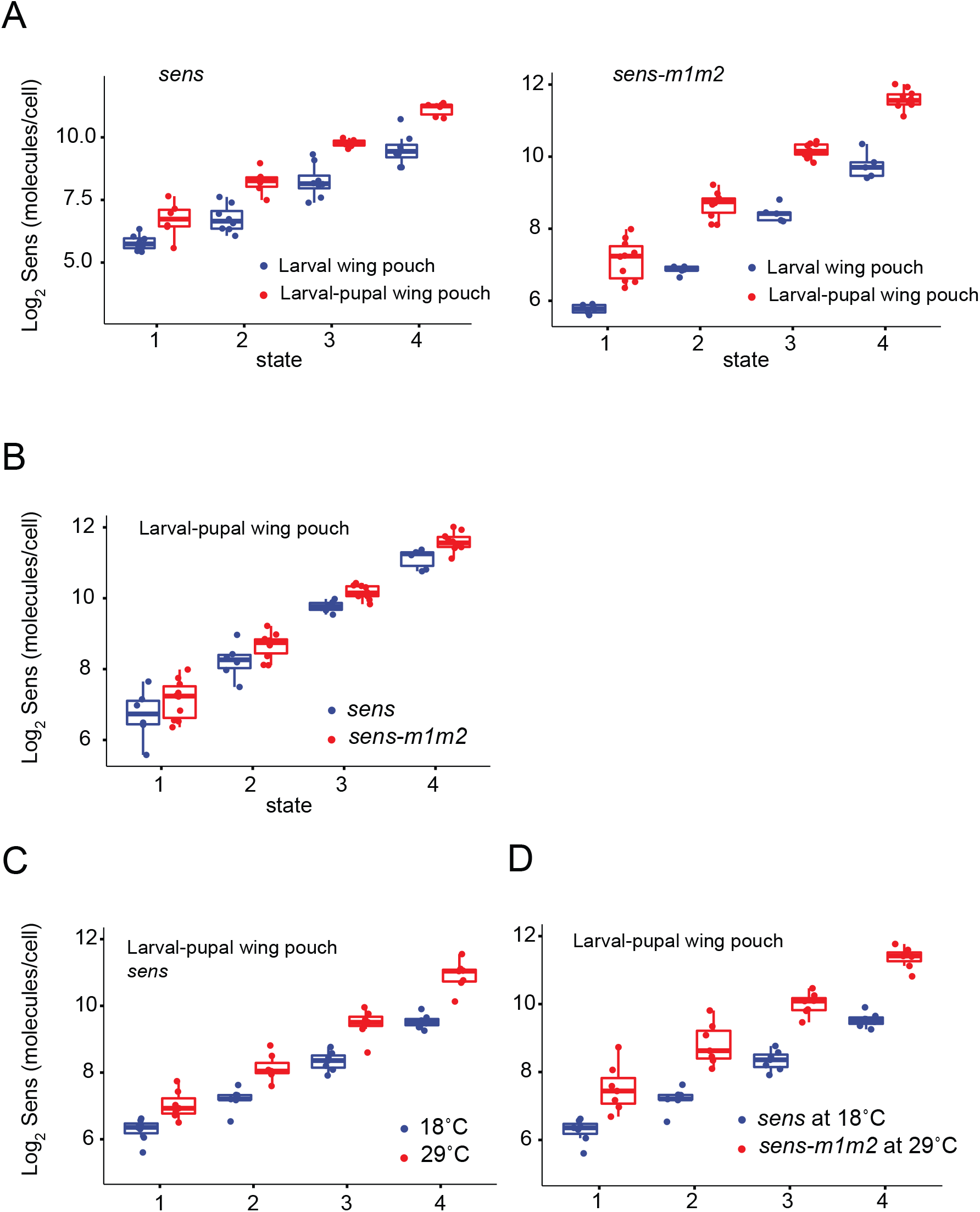
Sens states display log-linear increases irrespective of absolute abundance, related to Figures 5 and 6. Log2-scaled mean Sens protein numbers for cells in the four states are plotted for various experimental conditions. Each dot represents an estimate derived from one wing pouch sample. Multiple samples were analyze for each stage and condition. Boxplots show the median and interquartile range (25 – 75%), and whiskers represent the 1.5x interquartile range. **(A)** Wing pouches are from the larval (blue) and larval-pupal (red) stages of development for animals rescued with wild type sfGFP tagged *sens* (left) or miR-9a binding site mutant *sens-m1m2* (right). **(B)** Wing pouches are from larval-pupal animals rescued with *sens* (blue) or *sens-m1m2* (red). **(C)** Larval-pupal wing pouches expressing the wild type *sens* transgene grown at either 18°C (blue) or 29°C (red). **(D)** Comparison of larval-pupal wing pouches expressing the wild type *sens* transgene grown at 18°C (blue) or mutated *sens-m1m2* transgene grown at 29°C (red). See also Figure S7B.

**Figure S9.**
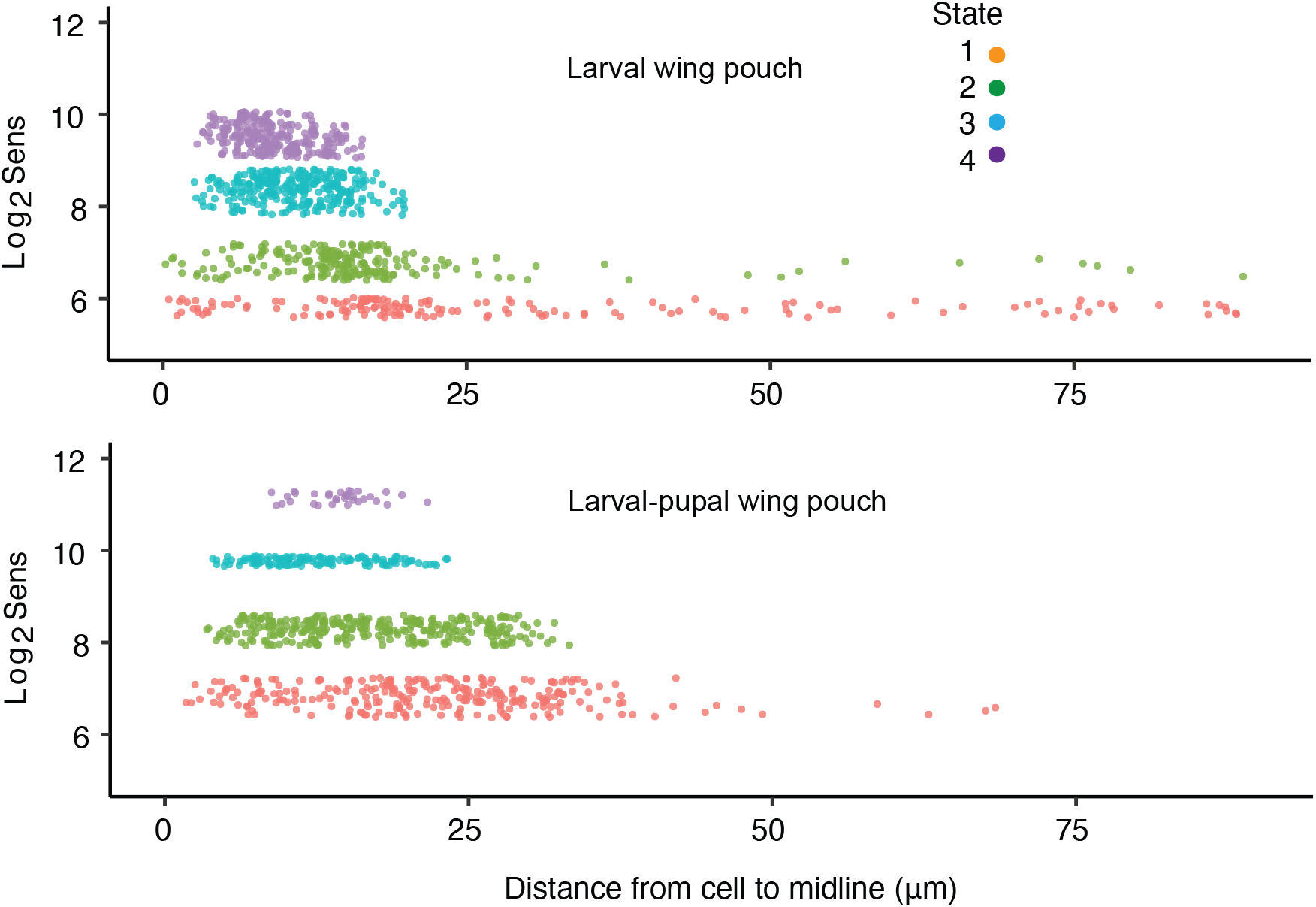
Spatial distribution of Log-scaled Sens expression, related to Figures 4 and 6. Single cells were classified into one of the states according to their Sens protein number relative to the mean Sens number in states. Cells are color coded according to state classification as shown and plotted as a function of their distance to the midline on the x-axis and log2-scaled Sens protein number on the y-axis. Shown are cells analyzed from one wing pouch taken at the larval stage (top) or larval-pupal transition stage (bottom).

